# mRNA-based influenza vaccine expands breadth of B cell response in humans

**DOI:** 10.1101/2024.10.10.617255

**Authors:** Hanover C. Matz, Tae-Geun Yu, Julian Q. Zhou, Lowrey Peyton, Anders Madsen, Fangjie Han, Aaron J. Schmitz, Stephen C. Horvath, Kritika Dixit, Hunter K. Keplinger, Benjamin S. Strnad, Mark J. Hoegger, William D. Middleton, Michael K. Klebert, Nina H. Lin, Raffael Nachbagauer, Robert Paris, Jackson S. Turner, Rachel M. Presti, Jiwon Lee, Ali H. Ellebedy

## Abstract

Eliciting broad and durable antibody responses against rapidly evolving pathogens like influenza viruses remains a formidable challenge^1,2^. The germinal center (GC) reaction enables the immune system to generate broad, high-affinity, and durable antibody responses to vaccination^3–5^. mRNA-based severe acute respiratory syndrome coronavirus 2 (SARS-CoV-2) vaccines induce persistent GC B cell responses in humans^6–9^. Whether an mRNA-based influenza vaccine could induce a superior GC response in humans compared to the conventional inactivated influenza virus vaccine remains unclear. We assessed B cell responses in peripheral blood and draining lymph nodes in cohorts receiving the inactivated or mRNA-based quadrivalent seasonal influenza vaccine. Participants receiving the mRNA-based vaccine produced more robust plasmablast responses and higher antibody titers to H1N1 and H3N2 influenza A viruses and comparable antibody titers against influenza B virus strains. Importantly, mRNA-based vaccination stimulated robust recall B cell responses characterized by sustained GC reactions that lasted at least 26 weeks post-vaccination in three of six participants analyzed. In addition to promoting the maturation of responding B cell clones, these sustained GC reactions resulted in enhanced engagement of low-frequency pre-existing memory B cells, expanding the landscape of vaccine-elicited B cell clones. This translated to expansion of the serological repertoire and increased breadth of serum antibody responses. These findings reveal an important role for the induction of persistent GC responses to influenza vaccination in humans to broaden the repertoire of vaccine-induced antibodies.

## Main text

Vaccines have reduced or eradicated the burden of many previously detrimental diseases^10^. However, for rapidly evolving pathogens such as seasonal influenza virus, it remains challenging to design fully efficacious vaccines that induce broadly neutralizing, durable antibody responses^1^. One component of the adaptive immune response to target for improved vaccine efficacy is the germinal center (GC) reaction. GCs are microanatomical structures that form in secondary lymphoid organs upon engagement of antigen and cognate B cells and T cells^3^, facilitate the Darwinian selection of high affinity antigen-specific B cell clones, and ultimately enhance antibody responses^4,5^. Additionally, GCs contribute to long-term protection by producing bone marrow plasma cells (BMPCs) and memory B cells (MBCs)^11^ which rapidly differentiate into antibody-secreting plasmablasts (PBs) upon antigen re-exposure^12^. Understanding how to design vaccines that effectively engage the GC reaction to promote both broadly binding antibodies and immune memory would greatly advance our ability to combat antigenically variable pathogens.

Influenza is a significant public health burden, causing 290,000-650,000 annual global deaths^2^. Seasonal influenza vaccines remain the most effective method for reducing the disease burden of influenza, with vaccines typically targeting the glycoprotein hemagglutinin (HA) utilized for viral entry into host cells^13^. However, due to antigenic drift and shift^14,15^, vaccines must be reformulated yearly^16^. Additionally, antigenic imprinting contributes to the generation of recall antibody responses that can be less effective against circulating viral strains^17,18^. These drawbacks underscore the need to determine how different vaccine platforms influence the GC response to influenza vaccination. The coronavirus disease 2019 (COVID-19) pandemic demonstrated that lipid nanoparticle-encapsidated messenger RNA (mRNA)-based vaccination is an effective alternative to conventional protein-based and inactivated virus-based vaccines in the context of a primary immune response^6,7^. However, influenza vaccination occurs in the context of secondary recall responses, and it is not known whether mRNA-based vaccination would produce a superior GC response to conventional vaccines. Furthermore, previous research has shown that both mRNA-based vaccines and inactivated virus-based vaccines can produce persistent GCs in humans, in some cases up to six months post vaccination^6,7,19^, but how these persistent GCs contribute to significant functional changes in the antibody repertoire remains to be fully determined.

We sought to characterize human B cell responses to an investigational mRNA-based quadrivalent seasonal influenza vaccine (mRNA-1010)^20^. We compared the dynamics of GC responses in vaccination cohorts receiving the 2022-2023 Northern Hemisphere inactivated quadrivalent influenza vaccine (Fluarix, n=15) or mRNA-1010 (n=14). Ultrasound-guided fine needle aspirations (FNAs) were used to directly sample the GC compartment in draining axillary lymph nodes.

### Robust humoral response to mRNA-based seasonal influenza vaccination

We conducted an observational study (WU397) of 29 healthy adults (ages 23-51, median age 33), with 15 participants receiving inactivated quadrivalent influenza vaccine (Fluarix) and 14 participants receiving mRNA-1010 encoding for the HA glycoproteins of the 2022-2023 Northern Hemisphere seasonal influenza virus strains (Extended Data Fig. 1a). Blood samples were collected at baseline and at 1, 2, 4, 8, 17, and 26 weeks post vaccination. A subset of these participants (n=11 Fluarix, n=6 mRNA-1010) enrolled for FNA specimens, collected at baseline and at 2, 8, 17, and 26 weeks post vaccination (Fig. 1a). The HA-specific PB responses in the blood were measured by enzyme-linked immune absorbent spot (ELISpot) assay against all four vaccine HA proteins. Frequencies of mean HA-specific IgG+ and IgA+ PBs per million peripheral blood mononuclear cells (PBMCs) peaked in both vaccine cohorts at 1 week post vaccination (Fig. 1b). At peak, participants receiving mRNA-1010 displayed significantly higher frequencies of IgG+ and IgA+ A/H1-specific and A/H3-specific PBs compared to participants who received Fluarix (Fig. 1c). IgM+ PB responses were greater in Fluarix participants (Extended Data Fig. 1b and c). No significant difference was observed in frequencies of influenza B IgG+ and IgA+ HA-specific PBs between the cohorts (Fig. 1c). Correspondingly, we observed significantly increased IgG plasma antibody titers to A/H1 and A/H3 as measured by ELISA as fold change over baseline at 4 weeks post vaccination in mRNA-1010 participants compared to Fluarix participants (Fig. 1d). For A/H3, these significantly higher titers persisted up to the final time point collected (week 17 or 26). No difference was observed between the cohorts for IgG serum titers to influenza B (Fig. 1d). In both cohorts, hemagglutination inhibition (HAI) titers against A/H1N1 and A/H3N2 significantly increased relative to baseline at 4 weeks post vaccination, but no significant difference was observed in peak titers between mRNA-1010 and Fluarix recipients despite the higher A/H3 titers measured by enzyme-linked immunosorbent assay (ELISA) in mRNA-1010 participants (Fig. 1e). We detected higher frequencies of circulating HA-specific MBCs by flow cytometric analysis (gating strategy Extended Data Fig. 1d) in the mRNA-1010 cohort at 4 weeks post vaccination and significant differences in the fold change of circulating MBCs at 4 and 17/26 weeks post vaccination over baseline (Extended Data Fig 1e and f). Overall, mRNA-1010 induced more robust antibody responses to A/H1 and A/H3 compared to inactivated virus vaccination.

**Fig. 1.**
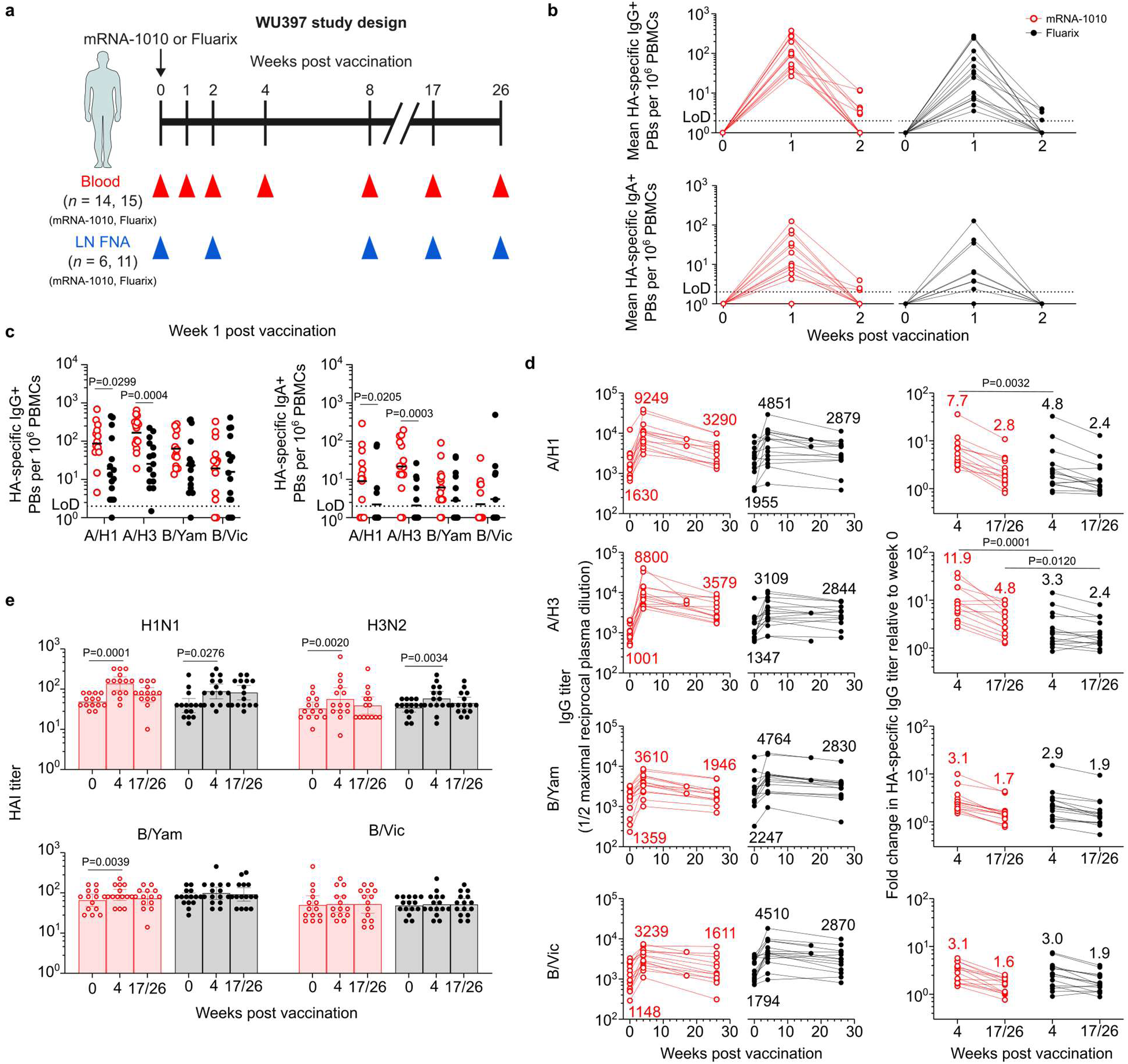
Robust antibody response to mRNA-based seasonal influenza vaccination. **a**, WU397 study design. We enrolled 29 healthy adults (ages 24-51) who received Fluarix (n=15) or mRNA-1010 (n=14) intramuscularly. Blood was collected before vaccination and at 1, 2, 4, 8, 17, and 26 weeks after vaccination. FNAs of ipsilateral axillary lymph nodes were collected before vaccination and at 2, 8, 17, and 26 weeks after vaccination. **b**, ELISpot quantification of mean HA-binding IgG- and IgA-secreting PBs in blood at baseline, 1, and 2 weeks after vaccination in mRNA-1010 (red) and Fluarix (black) participants. Numbers of HA-binding PBs were quantified against the four vaccine HAs and averaged. **c**, ELISpot quantification of HA-binding IgG- and IgA-secreting PBs at 1 week post vaccination in mRNA-1010 (red) and Fluarix (black) participants. Horizontal bars represent geometric means. *P* values determined by Mann-Whitney *U* test. **d**, Plasma IgG titers at baseline, 4, and 17/26 weeks post vaccination (left) and fold change in plasma IgG titers at 4 and 17/26 weeks over baseline (right) against the four vaccine HAs in mRNA-1010 (red) and Fluarix (black) participants. Numbers on left panels represent geometric mean titers for each time point; numbers on right panels represent mean fold changes. *P* values determined by Mann-Whitney *U* test. **e**, HAI titers for the four vaccine virus strains in mRNA-1010 (red) and Fluarix (black) participants at baseline, 4, and 17/26 weeks. Bars represent geometric mean with 95% confidence interval. *P* values determined by Wilcoxon matched pairs signed rank test. In **d-e**, for participants that did not complete a blood collection at week 26, samples from 17 weeks post vaccination were used for analysis (mRNA-1010, n=2; Fluarix, n=2).

### Germinal center response to mRNA-based vaccination

We next characterized the dynamics of GC responses in the mRNA-1010 and Fluarix participants. Both types of vaccine are injected into the deltoid muscle, which drains primarily to the lateral axillary lymph nodes. We used ultrasonography to identify and guide FNAs of accessible axillary lymph nodes on the side of immunization. The same lymph node was sampled in each participant at each time point. Flow cytometric analysis detected high proportions of HA-specific GC B cells in participants with active GC responses after vaccination with mRNA-1010 (representative results in Fig. 2a, gating strategy in Extended Data Fig. 2a). Frequencies of HA-specific GC B cells were higher in mRNA-1010 participants at 2 weeks post vaccination compared to Fluarix participants, and only in mRNA-1010 (3/6) participants did we detect HA-specific GC responses at 26 weeks (approximately 6 months) post vaccination (Fig 2b).

**Fig. 2.**
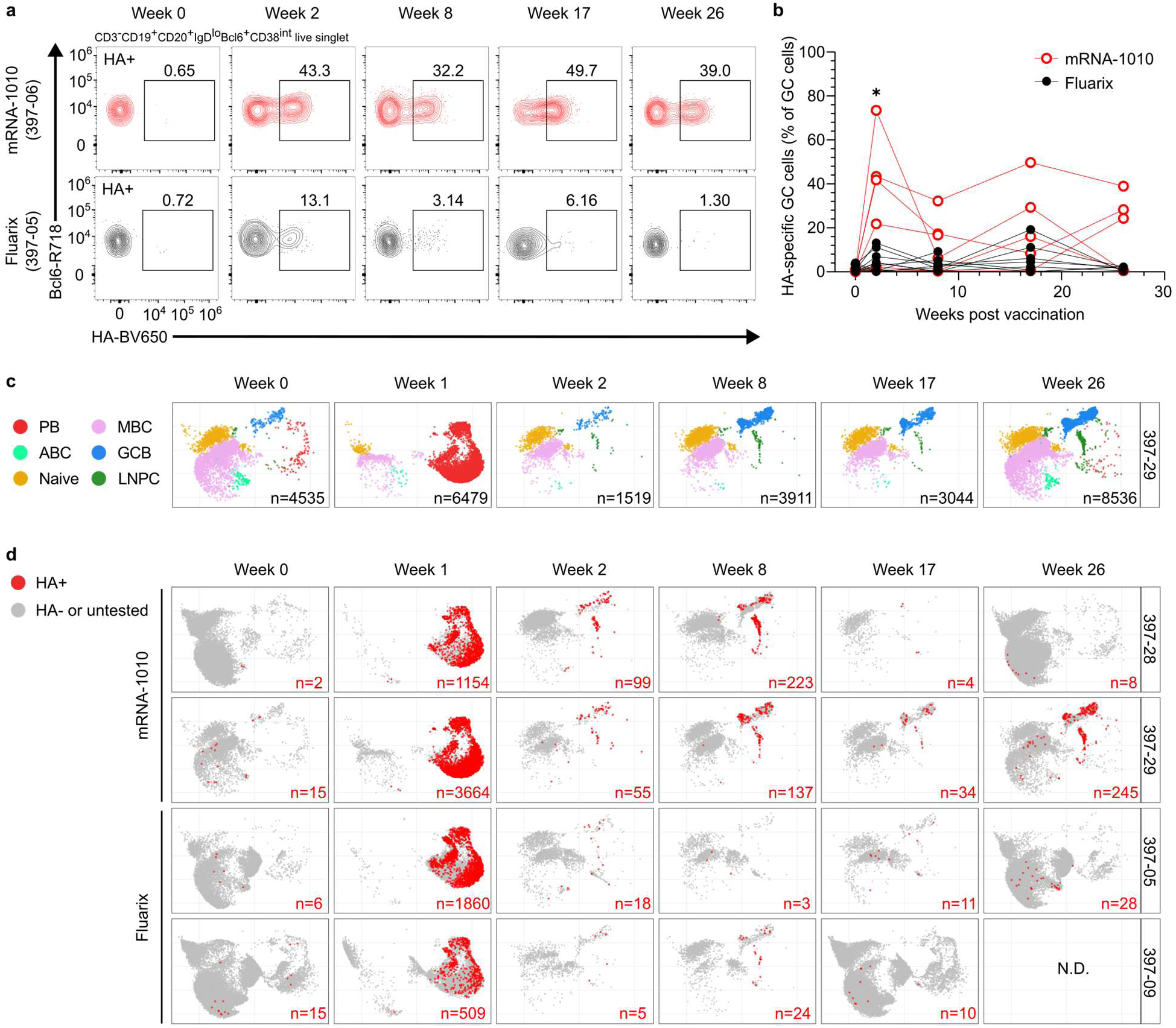
Characterizing the germinal center response to mRNA-1010. **a**, Representative flow cytometry plots of Bcl6 and HA-probe staining on CD3^-^CD19^+^CD20^+^IgD^lo^Bcl6^+^CD38^int^ live singlet lymphocytes in FNA samples at baseline, 2, 8, 17, and 26 weeks post vaccination. Top, representative mRNA-1010 participant (red, 06); bottom, representative Fluarix participant (black, 05). **b**, Frequencies of HA-specific GC B cells determined by flow cytometry in FNA samples from mRNA-1010 (red) and Fluarix (black) participants. *P* values determined by Mann-Whitney *U* test (*P=0.0365,* ^✱^*P*<0.05)**. c**, Representative uniform manifold approximation and projection (UMAP) plots of transcriptional clusters of B cells from baseline (sorted MBCs, FNA), 1 (sorted PBs), 2 (FNA), 8 (FNA), 17 (FNA), and 26 (sorted MBCs, FNA) weeks post vaccination (mRNA-1010 participant 29). Each dot represents a cell, colored by phenotype as defined by transcriptomic profile. Total numbers of cells are shown at the bottom right. PB, plasmablast; ABC, activated B cell; MBC, memory B cell; GCB, GC B cell; LNPC, lymph node plasma cell. **d,** UMAP plots of transcriptional clusters of B cells for all samples from mRNA-1010 (28, 29) and Fluarix (05, 09) participants as in **c**, with HA-specific clones as determined by mAb ELISA mapped onto transcriptional clusters. Numbers of HA-specific cells (red) are shown at the bottom right. Numbers of HA-specific clones for all participants are shown in Extended Data Table 1.

To further investigate the B cell response, single-cell RNA sequencing (scRNA-seq) analysis was performed on week 0 and week 17 or 26 (depending on sample availability) sorted total MBCs from PBMCs, week 1 sorted total PBs from PBMCs, and whole FNA samples from all time points for four mRNA-1010 participants (06, 17, 28, and 29) and three Fluarix participants (05, 09, and 22) with detectable GC responses (gating for MBC and PB sorting in Extended Data Fig. 2b). This allowed for identification of B cell subtypes (PB, GC, MBC etc.) based on transcriptomic profiles (Fig. 2c, Extended Data Fig. 3a-d, Supplementary Tables 1-6) and inference of B cell clonal relationships based on paired heavy and light chain BCRs. BCRs from the same clone shared heavy and light chain V and J genes and complementarity-determining region 3 (CDR3) lengths, as well as 85% similarity amongst the nucleotide sequences of their heavy chain CDR3s.

Clonally distinct BCRs from the PB, GC B cell, and LNPC compartment were expressed as mAbs (n=2989) and tested for HA-specificity by ELISAs (Extended Data Table 1). HA-specific B cell clones were visualized by overlaying on the B cell subtypes, with the majority of HA-specific BCRs derived from week 1 PBs (Fig. 2d and Extended Data Fig. 3e).

Assignment of HA-specificity to the scRNA-seq BCR data allowed us to analyze the clonal overlap between the GC and PB compartments in mRNA-1010 and Fluarix recipients. As the response to influenza vaccination is primarily a recall of HA-specific MBCs^21,22^, the week 1 PB compartment predominantly represents clones with prior antigen experience. B cell clones detected in both the GC (all time points) and week 1 PB compartment are thus most likely derived from MBCs. The majority of HA-specific GC B cell clones in all of the mRNA-1010 participants (median 87%) and one of the Fluarix participants (median 20%) overlapped with the week 1 PB compartment (Fig. 3a and Extended Data Table 2). However, a greater (albeit not statistically significant, *P*=0.0571, Mann-Whitney *U* test) proportion of the week 1 PB clones (median 14%) overlapped with the GC compartment in the mRNA-1010 cohort than the Fluarix cohort (median 2%), suggesting a wider pool of MBCs may have been engaged by the mRNA-1010 vaccine.

**Fig. 3.**
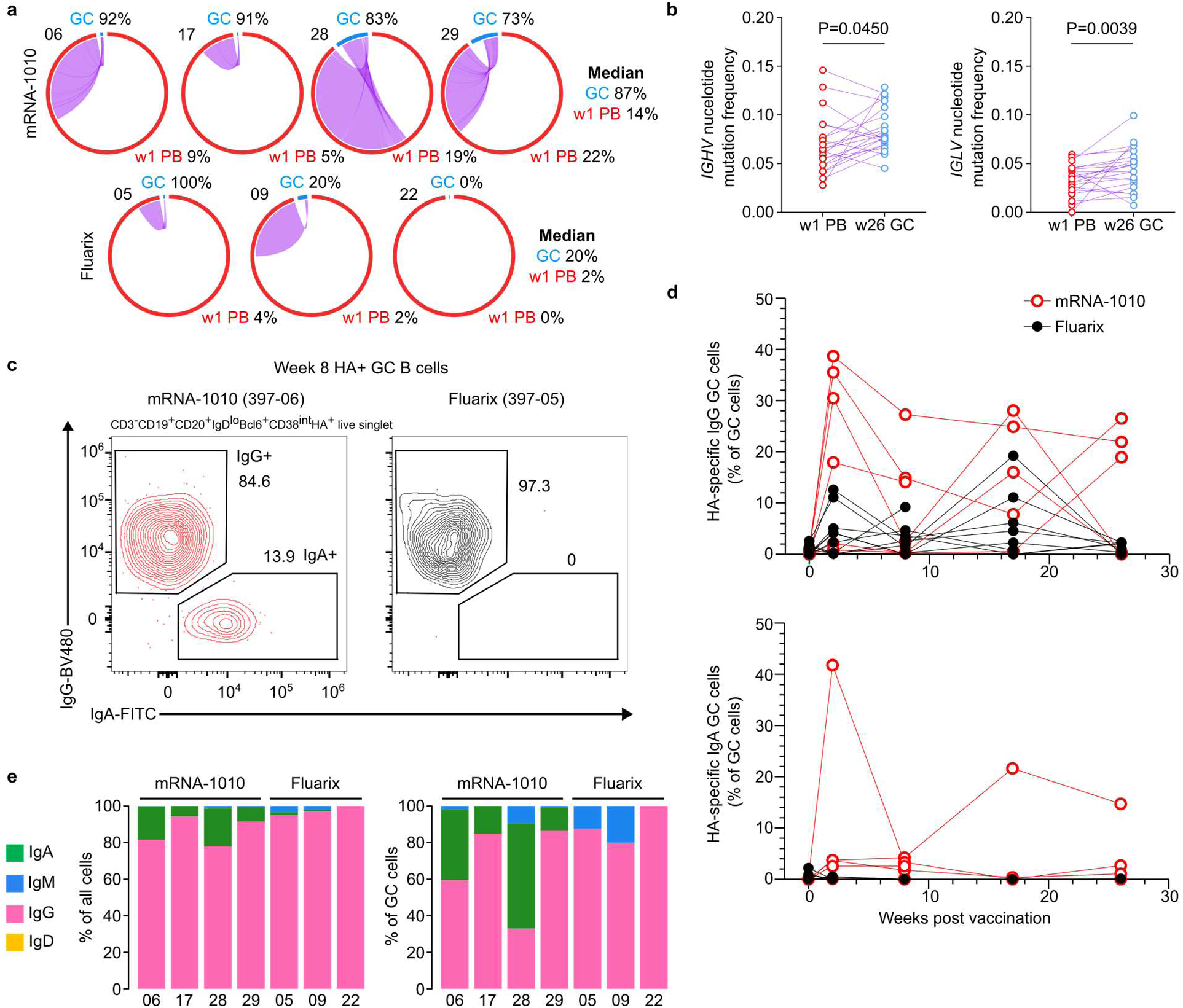
Germinal centers induced by mRNA-1010 recruit low frequency memory B cells and increase somatic hypermutation. **a**, Clonal overlap of sequences between PBs sorted from PBMCs 1 week after vaccination and GC B cells from all FNA time points among HA-specific clones for mRNA-1010 participants (06, 17, 28, 29, top) and Fluarix participants (05, 09, 22, bottom). Chord width corresponds to clonal population size; numbers of HA-specific clones are in Extended Data Table 2. Percentages are of GC B cell clones overlapping with PBs and PB clones overlapping with GC B cells. **b**, Nucleotide mutation frequency in the immunoglobulin heavy chain variable gene (IGHV) and light chain variable gene (IGLV) region for clonally related week 1 PBs and week 26 GC B cells (n=21) from mRNA-1010 participants. *P* values determined by Wilcoxon matched pairs signed rank test. **c**, Representative flow cytometry plots of IgG and IgA staining on CD3-CD19^+^CD20^+^IgD^lo^Bcl6^+^CD38^int^HA^+^ live singlet lymphocytes in FNA samples at 8 weeks post vaccination. Left, representative mRNA-1010 participant (red, 06); right, representative Fluarix participant (black, 05). **d**, Frequencies of IgG+ (top) and IgA+ (bottom) HA-specific GC B cells determined by flow cytometry in FNA samples from mRNA-1010 (red) and Fluarix (black) participants. **e**, Proportions of isotypes of HA-specific cells from scRNA-seq; BCR specificity determined by mAb ELISA and cells identified in transcriptional scRNA-seq clusters from mRNA-1010 (06, 17, 28, and 29) and Fluarix (05, 09, and 22) participants. Left, all B cells in scRNA-seq data; right, only GC B cells in scRNA-seq data.

To determine if the persistent GCs identified in mRNA-1010 participants altered SHM and affinity maturation in these HA-specific overlapping clones, we identified 21 pairs of clonally related BCRs present as week 1 PBs and week 26 GC B cells across 3 of the 4 mRNA-1010 participants in the scRNA-seq data (no such paired clones were detected in Fluarix participants due to the absence of detectable HA-specific week 26 GC B cells in this cohort). In these 21 paired clones, we observed significantly increased immunoglobulin heavy chain variable gene (IGHV) and light chain variable gene (IGLV) nucleotide mutation frequencies in the week 26 GC B cell compartment compared to the week 1 PB compartment, indicating these clones had undergone further somatic hypermutation (SHM) in GCs (Fig. 3b). However, we did not detect significant increases in their binding affinities by biolayer interferometry (BLI) or ELISA (Extended Data Fig. 4a and b).

We further examined the flow cytometry data for evidence that low frequency MBCs may be engaged in mRNA-1010-induced GC responses. Amongst HA-specific GC B cells in FNA samples, we observed detectable frequencies of both IgG+ and IgA+ cells in mRNA-1010 participants, whereas the corresponding populations in Fluarix participants were predominantly IgG+ (representative data Fig. 3c). Analysis of the kinetics of the GC responses demonstrated IgA+ HA-specific GC B cells were detectable in mRNA-1010 participants up to 26 weeks post vaccination (Fig. 3d). Consistent with this observation, IgA+ GC B cells were detected by scRNA-seq in mRNA-1010 participants but not Fluarix participants (Fig. 3e). Phylogenetic analysis of the IgA clones suggested they potentially arose from both IgA+ MBCs and IgG+ MBCs that had class switched to IgA (Extended Data Fig. 4c). As the ELISpot PB data demonstrated IgA+ HA-specific PBs displayed on average a 5-fold lower frequency compared to IgG+ HA-specific PBs (Fig. 1c), and other studies suggest IgA+ influenza-specific MBCs are lower frequency than IgG+ MBCs^22,23^, these results collectively suggest mRNA-1010 engages low frequency MBCs in GC responses.

### Functional antibody repertoire changes as a result of mRNA vaccination

To examine differences in the secreted antibody repertoires between the two vaccination cohorts, we performed high-resolution proteomic analysis of immunoglobulin (Ig-seq) coupled with high-throughput sequencing of transcripts encoding BCRs (BCR-seq)^24^ to quantitatively characterize the compositions of the serum antibody responses at the individual clonotype level (Fig 4a). A clonotype is defined as a group of heavy chain variable region (V_H_) sequences that share germline V and J segments and also exhibit greater than 90% amino acid identity in the heavy chain CDR3 (CDRH3)^25^. We focused on the serological IgG repertoire specific to the A/H3 component of the vaccine (A/Darwin/6/2021) as we observed the greatest difference in A/H3-specific serum binding titers between the two vaccine cohorts, and A/H3 is typically the most varied year-to-year vaccine component. From four mRNA-1010 participants (06, 17, 28, and 29) and four Fluarix participants (05, 09, 20, and 22) who (excluding 20) had robust GC responses detected by FNA, we isolated A/H3-specific serum IgG to delineate the serological repertoires (CDRH3 peptides) by mass spectrometry at baseline (week 0), peak (week 4), and final (week 17/26) time points. This was coupled with bulk BCR-seq from week 1 PBMCs to assign full length V_H_ sequences to clonotypes. We subsequently categorized individual IgG clonotypes as “pre-existing” or “vaccine-elicited” based on their presence or absence in serum, respectively, at baseline before vaccination (Fig. 4b and Extended Data Fig. 5a). We note that “vaccine-elicited” does not necessarily indicate arising from naïve responses but rather to induction of detectable IgG secretion following vaccination, which primarily arises from reactivated MBCs differentiating into antibody-secreting cells.

**Fig. 4.**
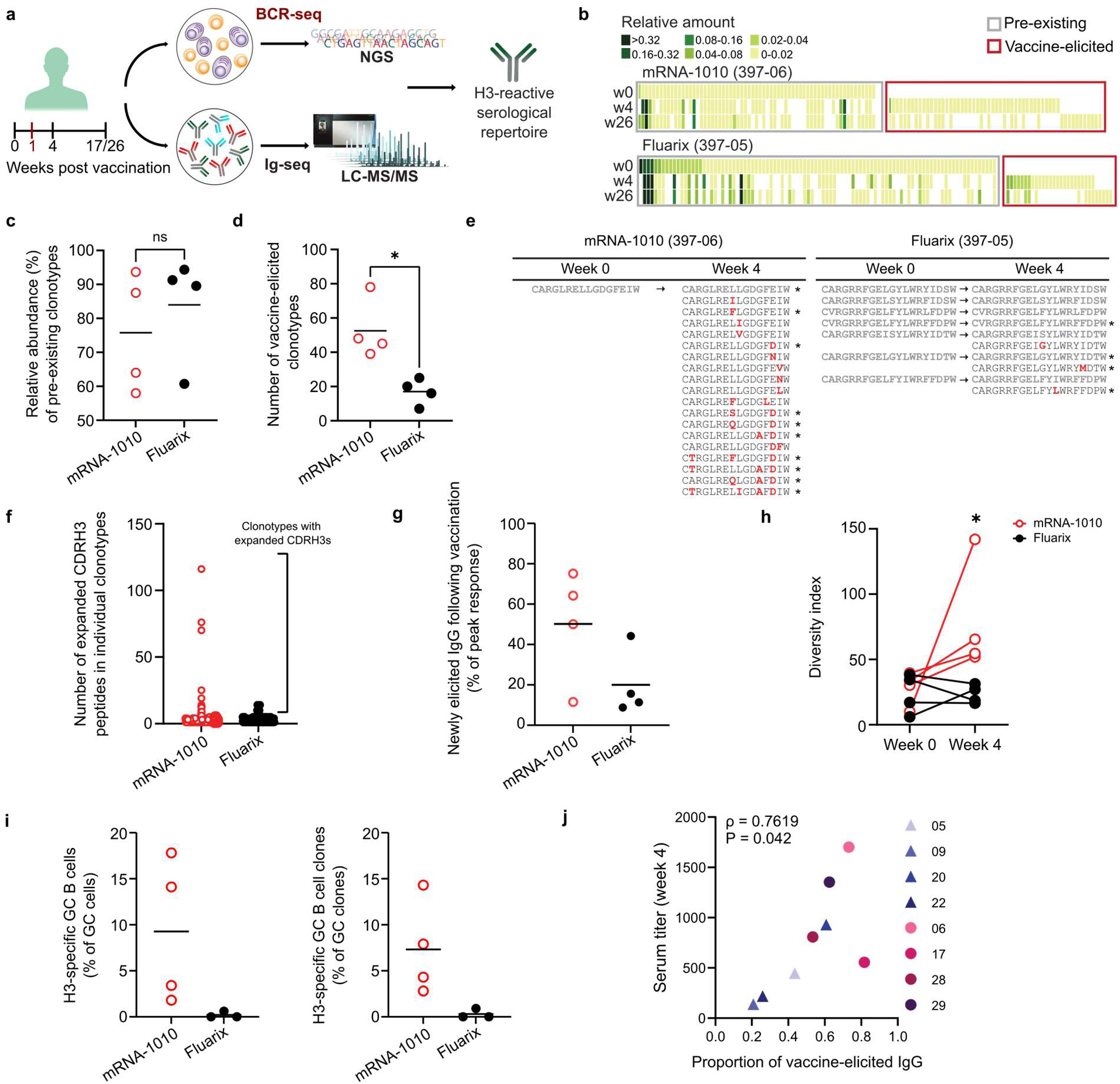
Delineation of H3-specific serological repertoires following vaccination. **a**, A schematic illustration of the proteomic analysis of serum immunoglobulins (Ig-seq) combined with high-throughput sequencing of B cell transcripts (BCR-seq) to identify the serological antibody repertoire to H3. **b**, Heat maps showing the relative amounts of IgG clonotypes comprising the serological repertoire against H3 at different time points from representative subjects (06, mRNA-1010; 05, Fluarix). Each column represents a unique serum IgG clonotype, with its relative amount determined through proteomic analysis. **c**, Relative abundance of pre-existing antibody clonotypes in serum at peak responses. Each data point represents an individual participant, quantified via Ig-seq. **d**, Number of serum vaccine-elicited H3-specific IgG clonotypes in serum at peak responses. Each data point represents an individual participant. **e**, Diversification of CDRH3 peptides detected sequences from representative clonotypes of the serological repertoire against H3 at baseline and peak time point. New amino acid replacement mutations are highlighted in red. Asterisks (*) indicate clones identified in scRNA-seq data in the week 1 PB compartment. **f**, Number of unique CDRH3 peptides that constitute individual IgG clonotypes. **g**, Relative abundance of newly elicited IgG following vaccination detected via Ig-seq. **h**, Change in diversity indices from baseline to peak response. Each line represents the change in diversity indices for an individual participant. **i**, Predominance of H3-specific GC cells determined by scRNA-seq data. Each data point indicates the fraction of H3-specific GC cells or GC clones for a specific participant. **j**, Correlation between serum titer at peak response and the proportion of vaccine-elicited IgG at peak response. ρ represents Spearman’s correlation coefficient. For **c**, **d**, **g**, and **i**, individual data points and the mean values are shown. For **c**, **d**, and **h,** statistical analyses were performed using two-tailed Mann-Whitney U test (^✱^*P*<0.05). For **j**, two-tailed Spearman rank correlation test was performed.

In the mRNA-1010 and Fluarix participants, on average, pre-existing antibody clonotypes comprised 75.8% (ranging from 58.0% to 93.6%) and 84.0% (ranging from 60.7% to 94.4%), respectively, of the total A/H3-specific serum IgG at peak responses (Fig. 4c), demonstrating the predominant contributions of boosted pre-existing clonotypes in the vaccine responses. While relative abundances of pre-existing clonotypes were not significantly different between the two vaccine cohorts, we noted that the number of vaccine-elicited serum IgG clonotypes at peak responses was significantly higher in mRNA-1010 participants (Fig. 4d). Furthermore, among highly abundant individual pre-existing clonotypes identified in serum at peak responses from the two vaccination cohorts, there were many clonotypes from mRNA-1010 participants drastically expanded following vaccination with increased diversity in the CDRH3 peptides within single IgG clonotypes (Fig. 4e and f). This was in comparison to the pre-existing clonotypes identified in Fluarix participants, which largely remained unchanged in CDRH3 peptide composition at week 4 (Fig. 4e and f). Several of these newly detected CDRH3 peptides within pre-existing clonotypes could be identified in the week 1 PB compartment of our scRNA-seq data (denoted by asterisk in Fig. 4e), suggesting that some of these peptides originated from restimulation of divergent MBCs that had previously undergone SHM. When accounting for such vaccine-induced diversification of CDRH3 giving rise to new IgG in serum, three of the mRNA-1010 participants (06, 17, and 29) exhibited high abundance (≥ 50%, mean 50.2% for all four participants) of newly elicited IgG in peak responses (Fig. 4g). In comparison, three of the Fluarix participants showed low abundance (<16%, mean 20% for all four participants) of new IgG contributing to the peak serum IgG responses. These differences between cohorts were reflected in the diversity index^26–28^ of the detected CDRH3 peptides; mRNA-1010 and Fluarix participants showed similar diversity indices at baseline but the diversity indices for mRNA-1010 participants were significantly higher at peak responses (Fig. 4h). Based on the scRNA-seq data, on average, 9.3% of GC B cells and 7.3% of GC B cell clones were H3-specific in mRNA-1010 participants, compared to 0.2% of GC B cells and 0.3% of GC B cell clones in Fluarix participants (Fig. 4i), suggesting H3-specific clones were more likely to be recruited into GCs in mRNA-1010 participants. We observed that the increased diversity of the vaccine-elicited serum IgG repertoires correlated with higher serum binding titers at peak responses across all the participants (Fig. 4j), and vaccine-induced expansion of the serological repertoires was maintained until the final time point (Extended Data Fig. 5b). These results suggest mRNA-1010 stimulated greater expansion of the H3-specific serological repertoires through both MBC recall into PB responses and recruitment into GCs for further SHM.

To further investigate the additional maturation of A/H3-specific serum IgG clonotypes in mRNA-1010 participants following vaccination, we performed B cell clonal lineage analysis on the most abundant A/H3-specific serum IgG clonotypes. In particular, we focused on three serum IgG clonotypes detected each from participants 06 and 28 that were among the most abundant serum clonotypes at peak responses (Fig. 5a and Extended Data Fig. 6a) and constructed phylogenetic trees based on week 1 bulk BCR-seq and scRNA-seq. These pre-existing clonotype lineages contained both pre-existing CDRH3 peptides detected at baseline and peak, as well as newly elicited serum IgG (as in Fig. 4e). Overall, we observed that vaccine-elicited serum IgG either mapped to branches maturing from pre-existing IgG or emerged as divergent and expanding branches. We also observed one or more CDRH3 peptides identified as clonally related GC B cells through scRNA-seq and located in branches corresponding to vaccine-elicited serum IgG in five of the six lineage trees (Fig. 5a, highlighted box, and Extended Data Fig. 6a). This data illustrates a subset of vaccine-elicited serum antibodies belong to lineages that are recruited into GCs following vaccination for further affinity maturation. In contrast, in Fluarix participants we rarely observed subbranches of lineage trees further maturing from pre-existing IgG, and no clonally related GC B cells were identified in the lineages we analyzed (Fig. 5a and Extended Data Fig. 6b). Thus, the lineage analysis confirms that mRNA-1010 likely stimulates divergent MBC clones that contribute to the serological repertoire and undergo further SHM in GCs.

**Fig. 5.**
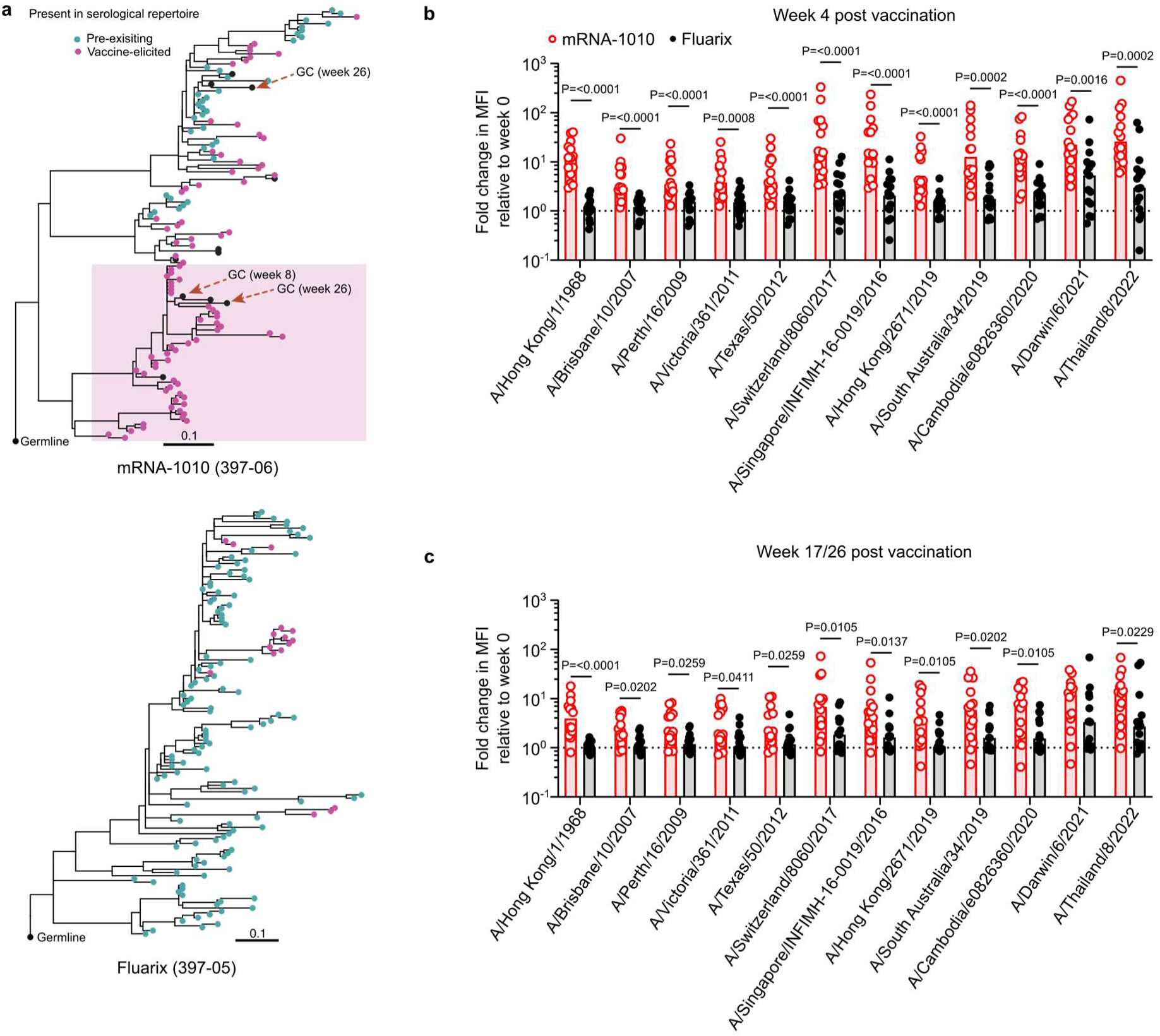
Increase in breadth of the H3-specific serological repertoire. **a**, B cell lineage trees of selected clonotypes. Each node is colored based on whether its CDRH3 sequence is categorized as pre-existing (turquoise) or vaccine-elicited (indigo) as determined by Ig-seq. Black color indicates CDRH3 sequences not detected in circulation. Branch lengths correspond to SHM per site, according to the scale bar. GC B cells are highlighted with the time points at which they were detected based on the scRNA-seq data. A sub-branch of BCRs with CDRH3 sequences matching to vaccine-elicited IgG and GC B cells is highlighted in a box. Top, mRNA-1010 participant (06); bottom, Fluarix partiicpant (05). **b**, Fold change in MFI of binding of plasma samples at 4 weeks post vaccination over baseline for mRNA-1010 (red) and Fluarix (black) participants to beads coated in a panel of A/H3 glycoproteins. **c**, Fold change in MFI of binding of plasma samples at 17/26 weeks over baseline for mRNA-1010 (red) and Fluarix (black) participants to beads coated in a panel of A/H3 glycoproteins. Samples collected at 17 weeks post vaccination were used for patients that did not complete a blood draw at 26 weeks (mRNA-1010, n=2, Fluarix, n=2). For **b** and **c**, *P* values were determined by Mann-Whitney U test. Bars represent median values.

To determine the impact of the vaccine-induced expansion of the serological repertoires on the functional capabilities of the antibody response, we measured the binding of final time point plasma samples against twelve antigenically diverse H3N2 influenza virus HA proteins covering more than 50 years of viral evolution, most of which were included in past recommended vaccines (Extended Data Fig. 7a). The higher variability of H3N2 compared to H1N1 or influenza B virus strains allowed us to test the binding breadth of antibody responses in the two vaccine cohorts. We tested plasma antibody binding to multiplex fluorescent beads coated with A/H3 protein from 12 H3N2 strains. Comparing fold change in median fluorescent intensity (MFI) over baseline (week 0), we observed significantly higher fold changes in plasma antibody binding to the antigenically divergent A/H3 strains in the mRNA-1010 cohort at peak (week 4) and final time points (week 17 or 26), with the most significantly higher binding for the oldest (and most divergent) strain, A/Hong Kong/1/1968 (Fig. 5b and c). Similar analysis conducted on H1N1 HA proteins also exhibited increased breadth of binding at peak but not final time point samples (Extended Data Fig. 7b and c). Collectively, our data demonstrate mRNA-1010 vaccination induces greater diversification of the serological repertoire which translates to the higher total serum binding titers and greater binding breadths against diverse influenza virus strains.

## Discussion

We observed that mRNA-based seasonal influenza vaccination is a robust alternative to conventional inactivated virus vaccines. In participants receiving mRNA-1010, vaccination resulted in higher antibody titers to H1N1 and H3N2 seasonal influenza viruses. We did not observe significant differences in antibody titers for influenza B viruses, but recent optimization of the mRNA-1010 vaccine has improved influenza B responses and demonstrated higher antibody titers compared to a currently licensed standard-dose flu vaccine^29^. While the antibody titers observed in this study did not result in significant differences in HAI between cohorts, our data suggests mRNA-1010 better engages a diverse pool of lower frequency MBCs. Clones diversified by previous SHM are recalled into the PB response, expanding the serological repertoire, and a subset of these clones are recruited into persistent GCs which further diversify their BCRs for future responses. Combined, these processes result in greater breadth of the influenza virus HA-specific antibodies. The mechanism underlying this outcome is still not fully elucidated. It is possible mRNA vaccines deliver greater amounts of antigen to the draining lymph nodes, resulting in both the stimulation of low frequency MBCs and prolonged antigen duration to drive GC persistence. It is also possible the self-adjuvanting properties of lipid nanoparticle mRNA vaccines^30^ robustly activate either professional antigen presenting cells and/or B cells for a more potent immune response. The form of antigen presented (i.e. membrane bound) as a result of mRNA-1010 vaccination may also contribute to increased valency of antigen and thus stimulation of low frequency and/or low affinity MBCs. Broadly, our results demonstrate that while there is a biological ceiling to antibody affinity with repeat antigen exposure, MBCs may re-engage in GCs for further SHM in order to diversify the antibody repertoire. Sustained GCs stimulated by mRNA-1010 vaccination potentially enhance this expansion of the MBC repertoire by increasing the opportunity for clones to undergo repeated rounds of mutation. This diversification may provide anticipatory mutations to combat rapidly evolving pathogens such as influenza virus^31,32^. The functional benefit of MBC lineages diversified by SHM is apparent based on our Ig-seq and serological binding breadth analyses. The presence of CDRH3 peptide-diversified pre-existing clonotypes and abundant newly detectable secreted vaccine-elicited clonotypes in the serological repertoire of mRNA-1010 but not Fluarix participants demonstrates that the diversification of the available pool of secreted antibodies likely contributes to the improved binding breadth observed in the responses of the mRNA-1010 cohort. Thus, it is possible that repeat doses over multiple influenza seasons of a vaccine such as mRNA-1010 may broaden the influenza-specific MBC repertoire via SHM in persistent GCs in contrast to conventional inactivated vaccines which, lacking broad MBC stimulation or sustained GCs, narrow the antigenic landscape of recall responses. As a vaccine that elicits both strong protection against antigenically drifting seasonal influenza strains as well as broad binding against divergent strains is recommended for the development of a universal influenza vaccine^33^, the results suggest that mRNA-based vaccines would greatly contribute to advancing this goal.

---

**Extended Data Fig. 1.**
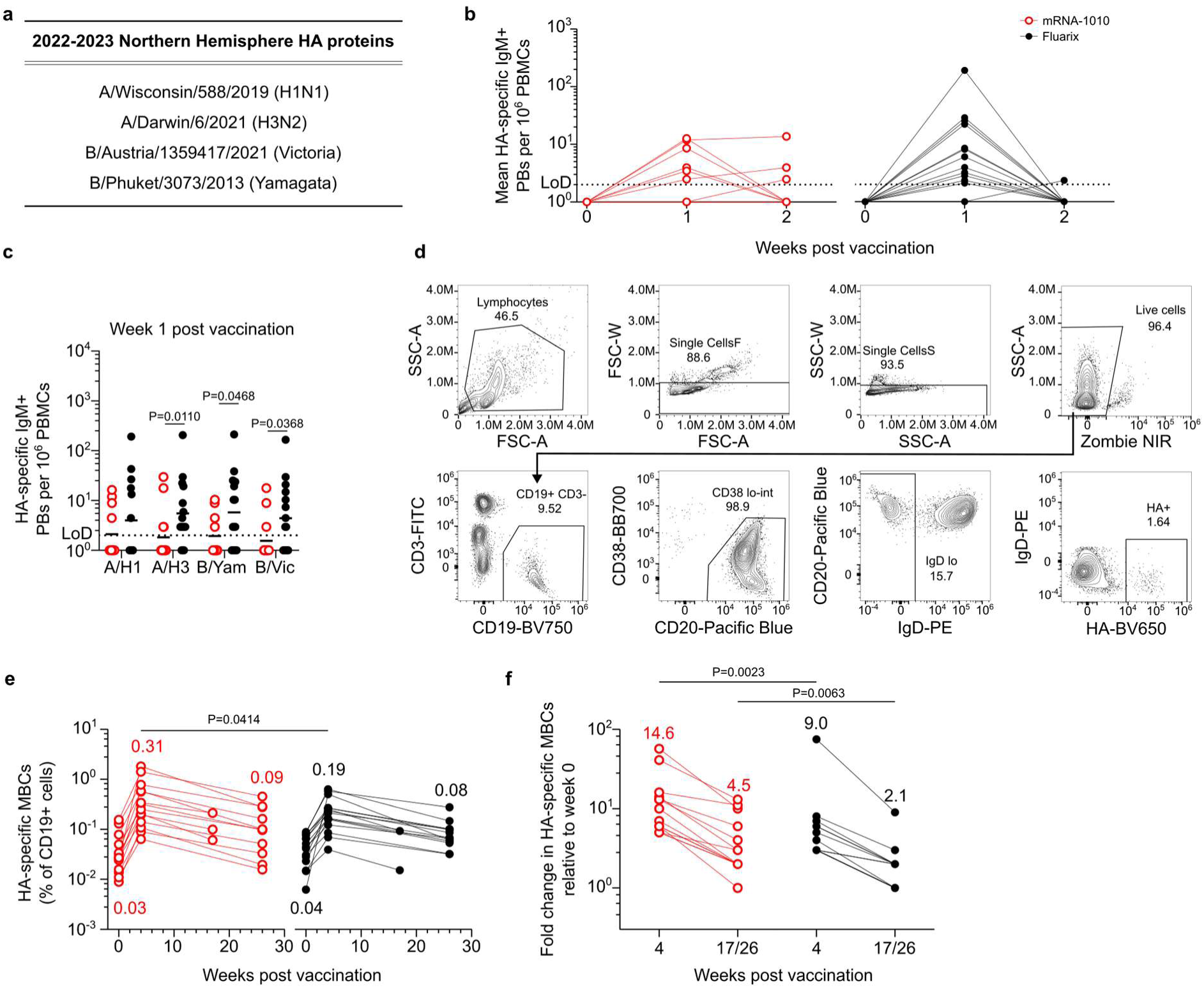
Humoral response to mRNA-1010. **a**, HA proteins encoded by mRNA-1010 for the 2022-2023 Northern Hemisphere influenza vaccine strains. Fluarix vaccine included the same strains, except H3N2 was egg-based (A/Darwin/9/2021). **b**, ELISpot quantification of mean HA-binding IgM-secreting PBs in blood at baseline, 1, and 2 weeks post vaccination in mRNA-1010 (red) and Fluarix (black) participants. Numbers of HA-specific PBs were quantified against the four vaccine HAs and averaged. **c**, ELISpot quantification of HA-binding IgM-secreting PBs at 1 week post vaccination in mRNA-1010 (red) and Fluarix (black) participants. Horizontal bars represent geometric means. *P* values determined by Mann-Whitney *U* test. **d**, Flow cytometry gating strategy for HA-specific MBCs from PBMCs **e**, Quantification of HA-specific MBCs as a percentage of CD19+ cells in blood by flow cytometry at baseline, 4, and 17/26 weeks post vaccination in mRNA-1010 (red) and Fluarix (black) participants. Numbers represent geometric mean frequencies of MBCs for each time point. **f**, Fold change in HA-specific MBCs as a percentage of CD19+ cells in blood by flow cytometry at indicated time points over week 0 for mRNA-1010 (red) and Fluarix (black) participants. Numbers represent mean values. In **e** and **f**, *P* values determined by Mann-Whitney *U* test. For participants that did not complete a blood collection at week 26 or PBMCs were not available, samples from 17 weeks post vaccination were used for analysis (mRNA-1010, n=3; Fluarix, n=2).

**Extended Data Fig. 2.**
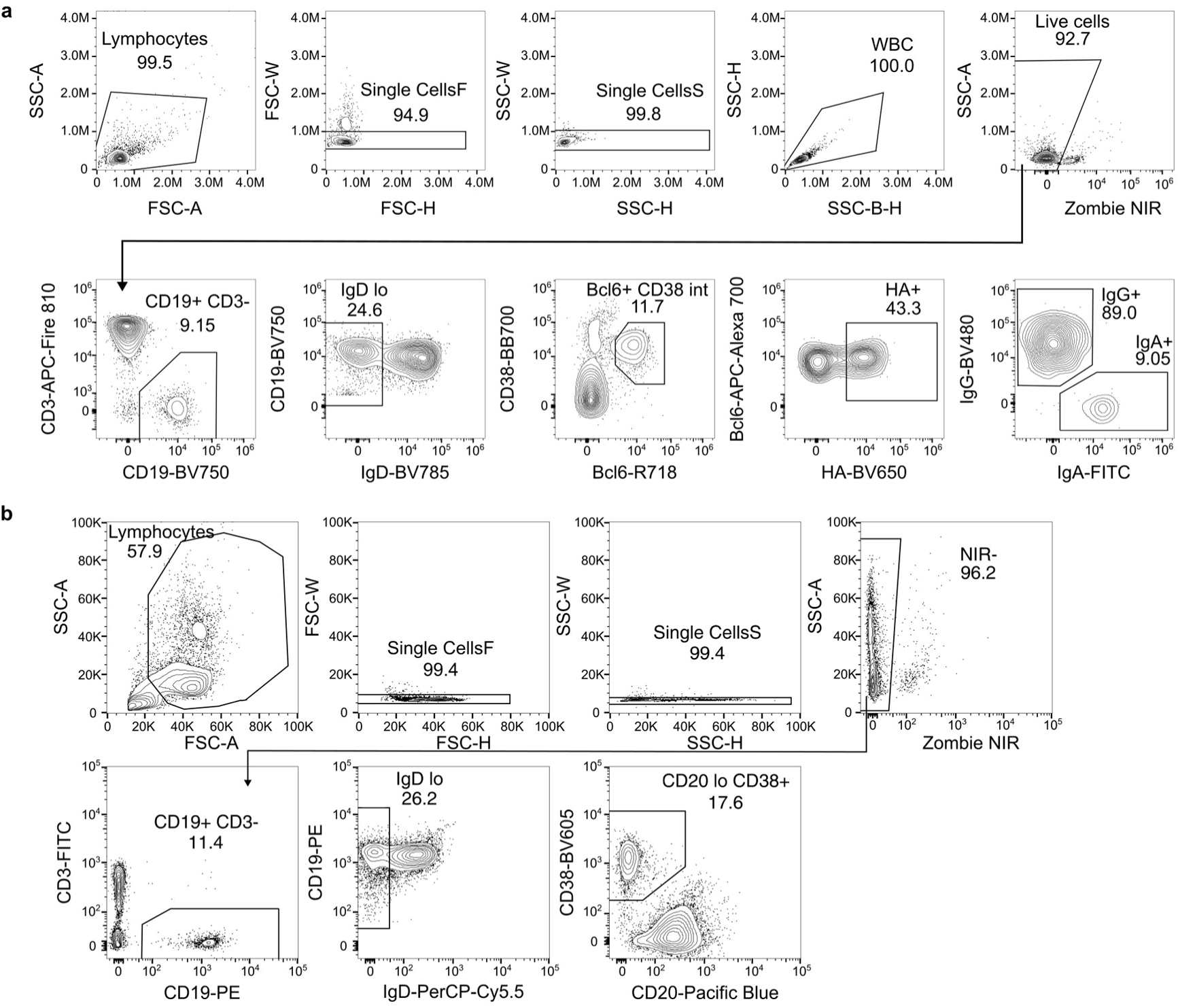
Flow cytometry gating strategies for analysis of B cell responses in vaccinees. **a**, Flow cytometry gating strategy for HA-specific GC B cells from FNAs. **b**, Gating strategy for sorting PBs and MBCs from peripheral blood. PBs were sorted as CD3^-^ CD19^+^IgD^lo^CD20^lo^CD38^+^ live singlet lymphocytes; MBCs were sorted as CD3^-^CD19^+^IgD^lo^ live singlet lymphocytes.

**Extended Data Fig. 3.**
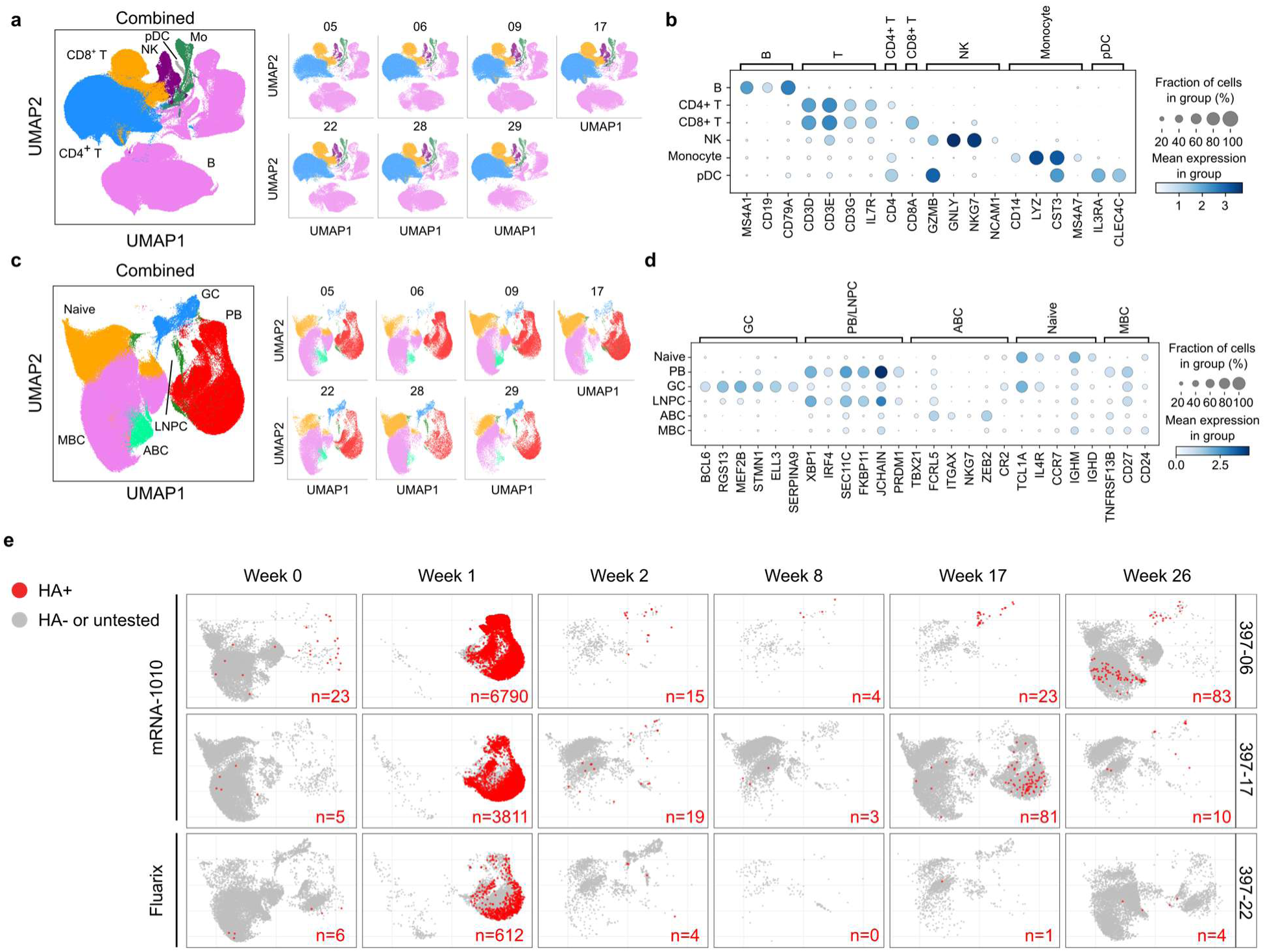
Identification of HA-specific B cell clones in lymph nodes and blood. **a** and **c,** UMAPs showing scRNA-seq transcriptional clusters of total cells (**a**) and of B cells (**c**) from PBs and MBCs sorted from blood and FNA of draining axillary lymph nodes combined. **b** and **d**, Dot plots for the marker genes used for identifying annotated clusters in **a** and **c**, respectively. **e**, UMAP plots of transcriptional clusters of B cells for all samples from mRNA-1010 (06, 17) and Fluarix (22) participant with HA-specific clones as determined by mAb ELISA mapped onto transcriptional clusters as in Fig. 2c. Numbers of HA-specific cells (red) are shown at the bottom right. Numbers of HA-specific clones for all participants are shown in Extended Data Table 1.

**Extended Data Fig. 4.**
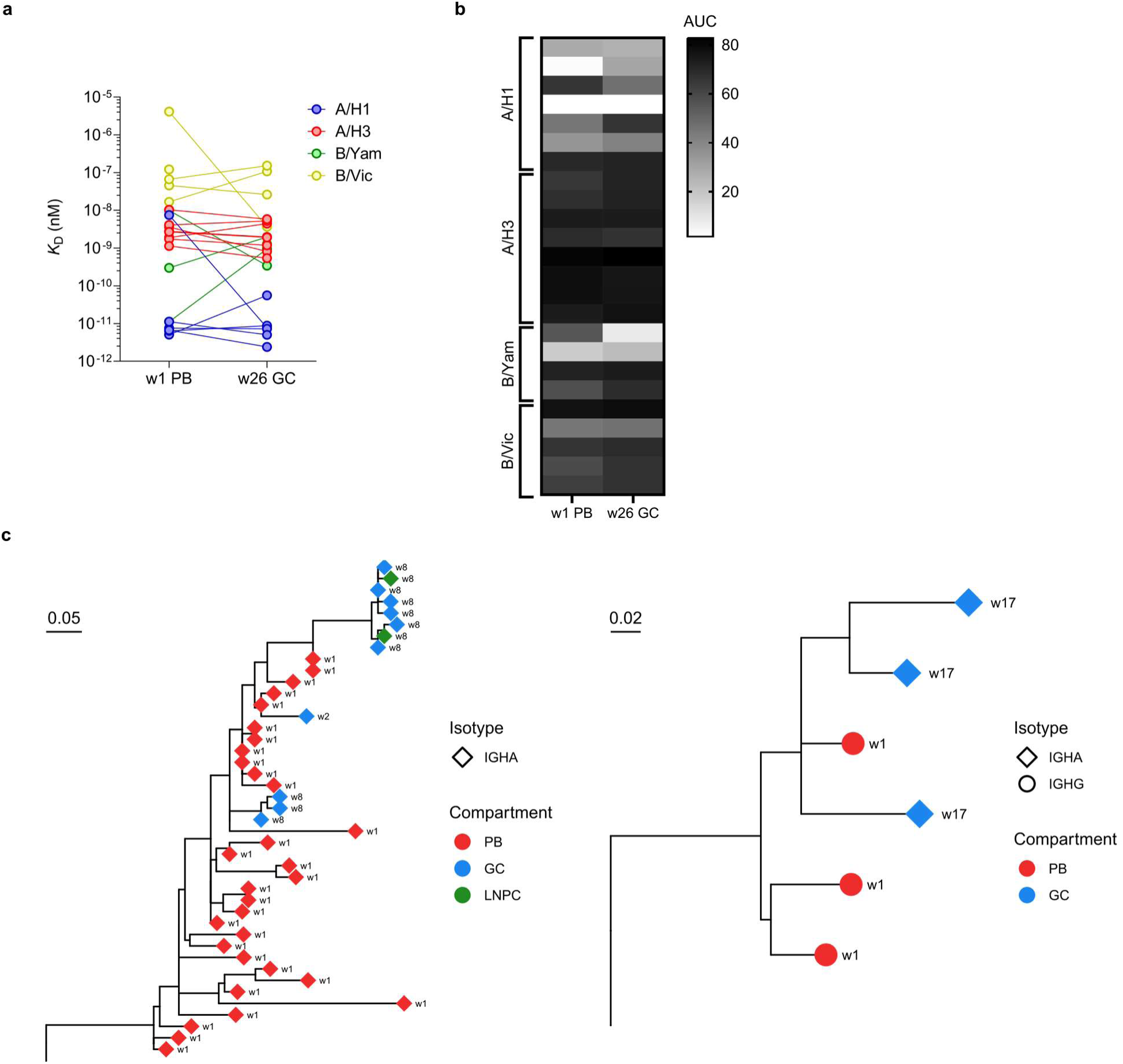
Analysis of germinal center antibody clones from mRNA-1010 participants. **a**, Equilibrium dissociation constant (*K*_D_) of Fabs (n=21) derived from paired week 1 PB and week 26 GC B cell clones of mRNA-1010 participants interacting with immobilized HA protein measured by BLI. Clones were tested against HA strains based on predetermined binding specificity by mAb ELISA; clones with multiple binding specificities were tested individually against each HA strain. **b**, Area under the curve (AUC) values for mAbs (n=21) derived from paired week 1 PB and week 26 GC B cell clones of mRNA-1010 participants as measured by binding ELISA against HA protein. Clones were tested against HA strains based on predetermined binding specificity by mAb ELISA; clones with multiple binding specificities were tested individually against each HA strain. **c**, Phylogenetic trees of representative HA-specific IgA GC B cell clones from scRNA-seq data of mRNA-1010 participants. Left (397-29) represents a clone likely derived from an IgA+ MBC; right (397-06) represents a clone likely derived from an IgG+ MBC that class switched to IgA. IGHA, immunoglobulin heavy chain alpha constant region; IGHG, immunoglobulin heavy chain gamma constant region. PB, plasmablast; MBC, memory B cell; GC, germinal center B cell; LNPC, lymph node plasma cell.

**Extended Data Fig. 5.**
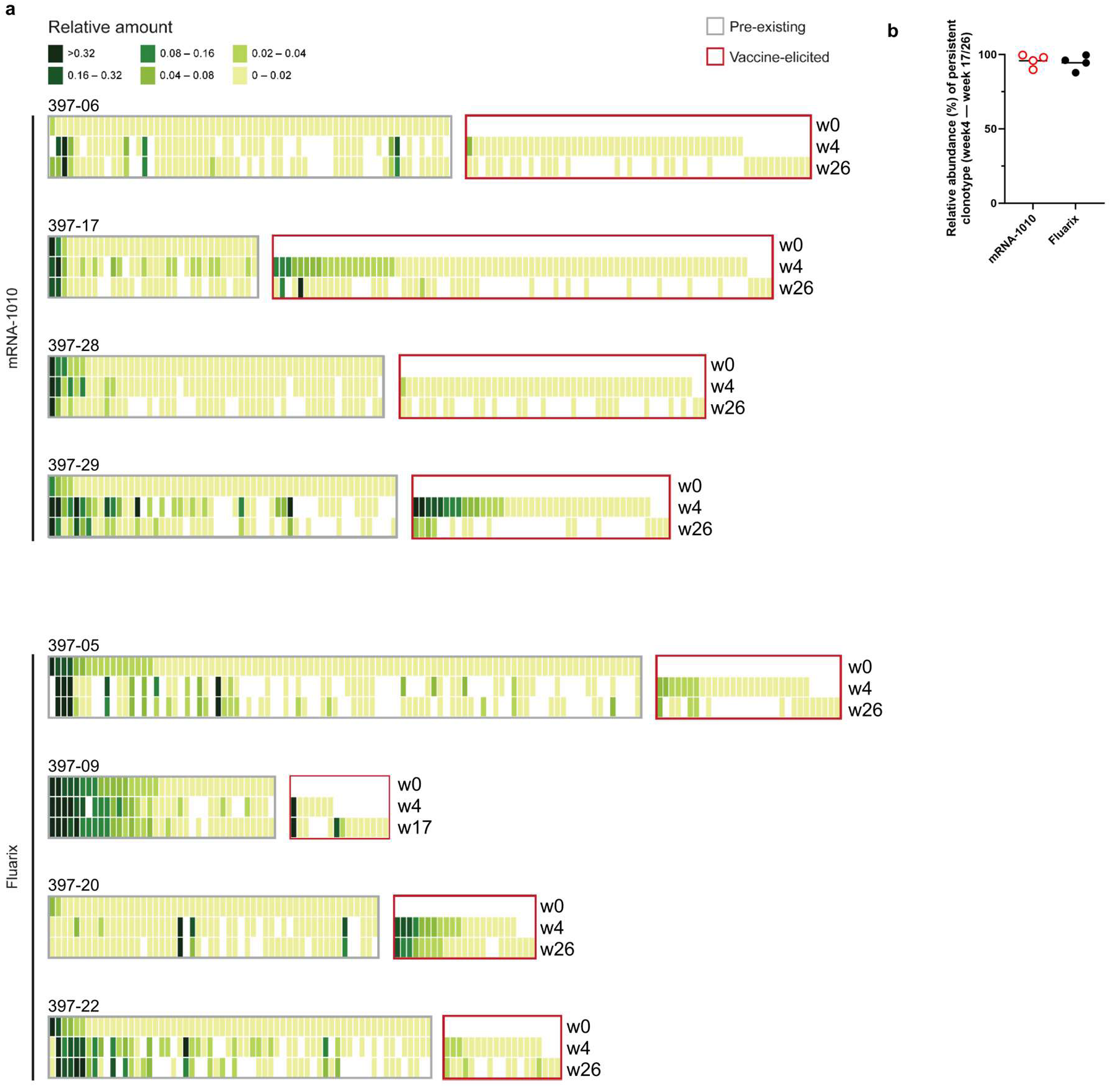
Antibody clonotypes comprising the H3-specific serological repertoire. **a**, Heat maps showing the relative amounts of IgG clonotypes comprising the serological repertoire against H3 at different time points from all participants. Each column represents a unique clonotype, with its relative amount determined through proteomic analysis. **b**, Persistence of post-vaccination serological repertoire between week 4 and week 17/26. Each data point represents the relative abundance of the serum IgG clonotype, quantified through Ig-seq, detected at week 17/26 which originate from week 4.

**Extended Data Fig. 6.**
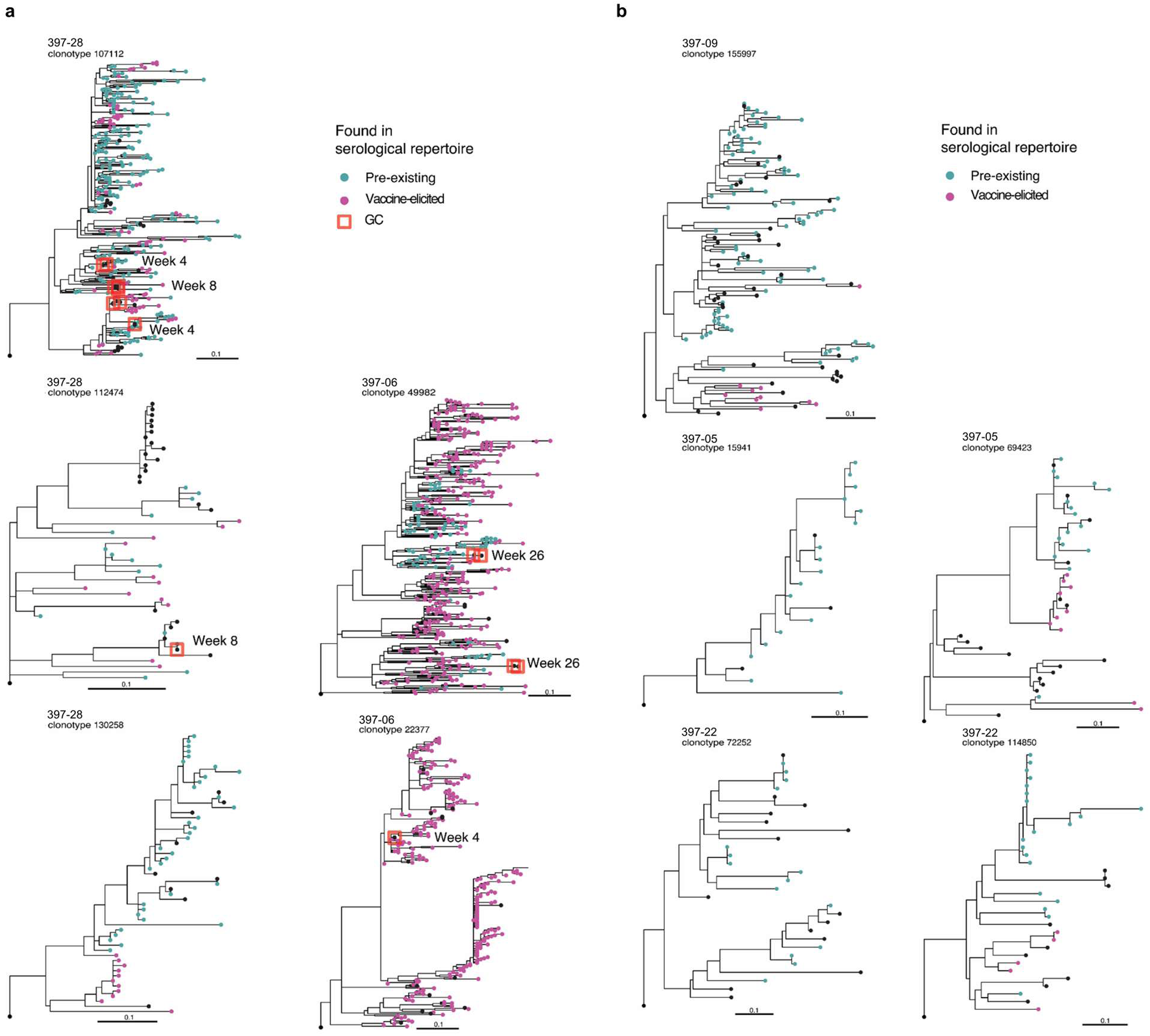
The B cell lineage tree of representative clonotypes identified from mRNA-1010 and Fluarix participants. **a** and **b**, B cell lineage trees of clonotypes abundantly present in serum identified from mRNA-1010 participants 06 and 28 (**a**) and Fluarix participants 05, 09, and 22 (**b**) are shown. Each node is colored based on whether its CDRH3 sequence is categorized as pre-existing (turquoise) or vaccine-elicited (indigo) as determined by Ig-seq. Black color indicates CDRH3 sequences not detected in circulation. Branch lengths correspond to SHM per site, according to the scale bar. GC B cells are highlighted with the time points at which they were detected based on the scRNA-seq data.

**Extended Data Fig. 7.**
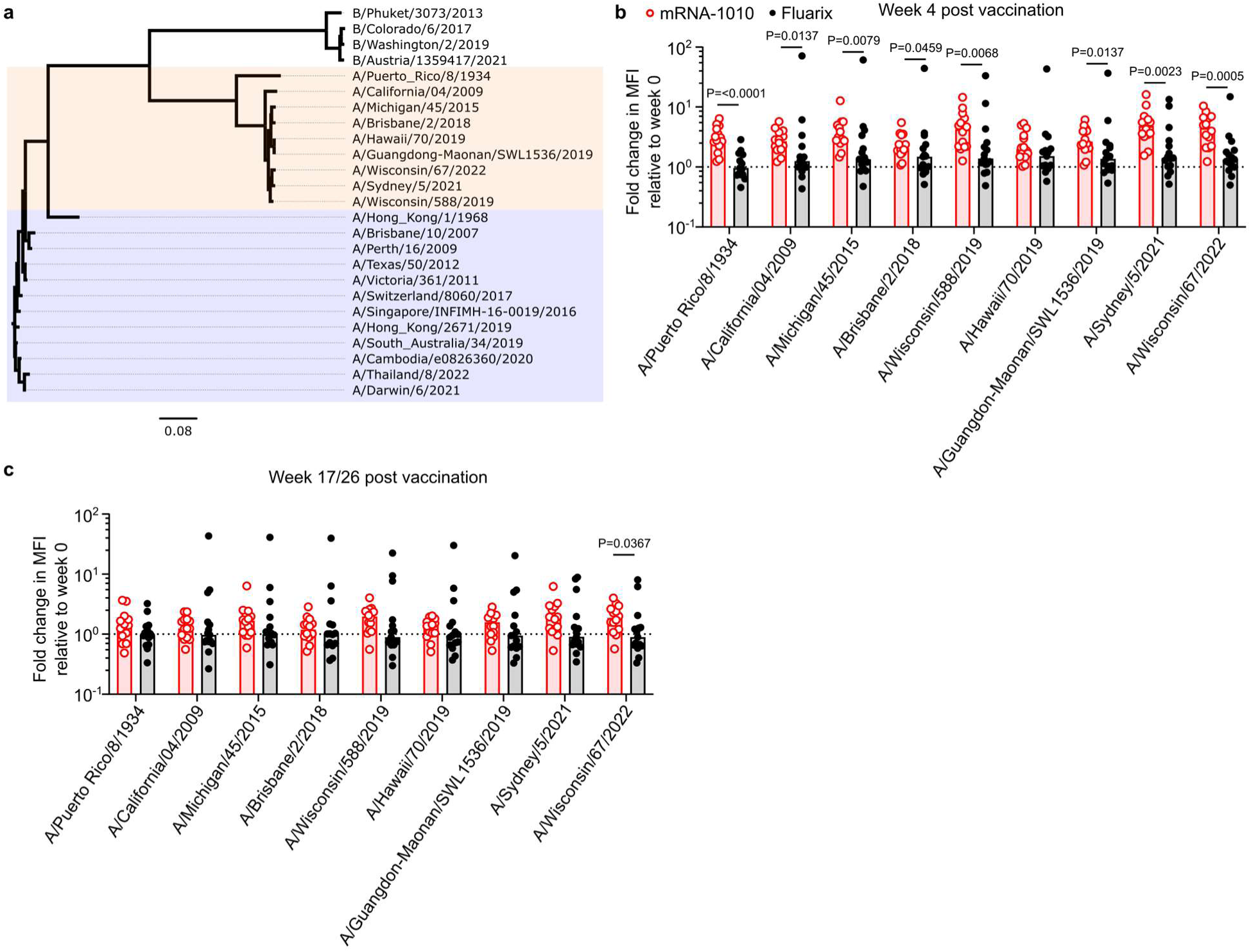
Breadth of binding of H1-specific serological repertoire. **a**, Phylogenetic tree of influenza strains. H1N1 strains used for bead assay are highlighted in orange; H3N2 strains are highlighted in blue. **b**, Fold change in MFI of binding of plasma samples at 4 weeks over baseline for mRNA-1010 (red) and Fluarix (black) participants to beads coated in A/H1 glycoproteins. **c**, Fold change in MFI of binding of plasma samples at 17/26 weeks over baseline for mRNA-1010 (red) and Fluarix (black) participants to beads coated in A/H1 glycoproteins. Samples collected at 17 weeks post vaccination were used for patients that did not complete a blood draw at 26 weeks (mRNA-1010, n=2, Fluarix, n=2). For **b** and **c,** *P* values were determined by Mann-Whitney *U* test; bars represent median values.

**Extended Data Table 1.**

| Monoclonal antibodies (mAbs) from WU397 participants |  |  |  |  |  |  |  |
| --- | --- | --- | --- | --- | --- | --- | --- |
| Participant | Vaccine | mAbs expressed | HA+ | H1+ | H3+ | B/Yam+ | B/Vic+ |
| 397-06 | mRNA-1010 | 439 | 259 | 126 | 89 | 39 | 46 |
| 397-17 | mRNA-1010 | 411 | 215 | 53 | 114 | 41 | 33 |
| 397-28 | mRNA-1010 | 174 | 81 | 26 | 32 | 25 | 16 |
| 397-29 | mRNA-1010 | 569 | 229 | 70 | 126 | 27 | 40 |
| 397-05 | Fluarix | 412 | 165 | 66 | 29 | 65 | 67 |
| 397-09 | Fluarix | 493 | 54 | 21 | 6 | 22 | 18 |
| 397-22 | Fluarix | 491 | 38 | 12 | 10 | 8 | 13 |
| Total |  | 2989 | 1041 | 374 | 406 | 227 | 233 |

Numbers of mAbs expressed from Fluarix (highlighted grey) and mRNA-1010 (highlighted red) participants. Total numbers of mAbs determined by ELISA to bind HA and numbers of mAbs determined to bind each vaccine HA strain are shown.

**Extended Data Table 2.**

| Participant | Vaccine | Week 1<br>PB clones | GC<br>clones | Clones in week 1 PB<br>and GC | % overlap<br>GC | % overlap<br>week 1 PB |
| --- | --- | --- | --- | --- | --- | --- |
| 397-06 | mRNA-1010 | 257 | 25 | 23 | 92.0 | 8.9 |
| 397-17 | mRNA-1010 | 213 | 11 | 10 | 90.9 | 4.7 |
| 397-28 | mRNA-1010 | 78 | 18 | 15 | 83.3 | 19.2 |
| 397-29 | mRNA-1010 | 207 | 63 | 46 | 73.0 | 22.2 |
|  |  |  |  | Median | 87.1 | 14.1 |
| 397-05 | Fluarix | 167 | 7 | 7 | 100.0 | 4.2 |
| 397-09 | Fluarix | 50 | 5 | 1 | 20.0 | 2.0 |
| 397-22 | Fluarix | 37 | 1 | 0 | 0.0 | 0.0 |
|  |  |  |  | Median | 20.0 | 2.0 |

Numbers of mAbs expressed from Fluarix (highlighted grey) and mRNA-1010 (highlighted red) participants determined to be HA-binding by ELISA with BCRs found in sorted PB clones at 1 week post vaccination and GC B cell clones.

**Supplementary Table 1.**

| Participant | Time Point | Tissue | Sorting | Replicate | BCR |  | 5' gene expression |  |  |  |
| --- | --- | --- | --- | --- | --- | --- | --- | --- | --- | --- |
|  |  |  |  |  | Pre-QC number of cells | Post-QC number of cells | Pre-QC number of cells | Post-QC number of cells | Median number of UMIs per cell | Median number of genes per cell |
| 397-05 | d0 | blood | IgDlo | 1 | 2788 | 2448 | 3263 | 3237 | 4170 | 1506 |
| 397-05 | d0 | blood | IgDlo | 2 | 3191 | 2779 | 3718 | 3680 | 4172 | 1508 |
| 397-05 | d0 | LN | NS | 1 | 514 | 458 | 1558 | 1391 | 3278 | 1453 |
| 397-05 | d0 | LN | NS | 2 | 523 | 479 | 1659 | 1477 | 3443 | 1505 |
| 397-05 | d8 | blood | PB | 1 | 4369 | 3758 | 5409 | 4839 | 5080 | 1799 |
| 397-05 | d8 | blood | PB | 2 | 4139 | 3647 | 5259 | 4738 | 5783.5 | 1829.5 |
| 397-05 | d15 | LN | NS | 1 | 935 | 855 | 4229 | 4013 | 4018 | 1627 |
| 397-05 | d15 | LN | NS | 2 | 1505 | 1379 | 5206 | 4961 | 4079 | 1631 |
| 397-05 | d57 | LN | NS | 1 | 706 | 647 | 2581 | 2480 | 3646 | 1499 |
| 397-05 | d57 | LN | NS | 2 | 798 | 706 | 2976 | 2891 | 3500 | 1447 |
| 397-05 | d121 | LN | NS | 1 | 1014 | 900 | 3493 | 3345 | 3782 | 1548 |
| 397-05 | d121 | LN | NS | 2 | 1254 | 1139 | 4102 | 3934 | 3852 | 1562 |
| 397-05 | d180 | blood | IgDlo | 1 | 6143 | 4844 | 5958 | 5929 | 4237 | 1517 |
| 397-05 | d180 | blood | IgDlo | 2 | 6838 | 5342 | 7045 | 7016 | 4147 | 1476 |
| 397-05 | d181 | LN | NS | 1 | 450 | 406 | 2998 | 1672 | 3811 | 1543 |
| 397-05 | d181 | LN | NS | 2 | 404 | 358 | 2706 | 1417 | 3877 | 1582 |
| 397-06 | d0 | blood | IgDlo | 1 | 9544 | 8085 | 9071 | 8952 | 4768 | 1596 |
| 397-06 | d0 | blood | IgDlo | 2 | 7594 | 6477 | 7292 | 7183 | 4895 | 1578 |
| 397-06 | d0 | LN | NS | 1 | 251 | 234 | 6298 | 6075 | 3492 | 1431 |
| 397-06 | d0 | LN | NS | 2 | 273 | 244 | 5313 | 5122 | 3411.5 | 1391 |
| 397-06 | d8 | blood | PB | 1 | 6258 | 5692 | 7956 | 6937 | 11145 | 2311 |
| 397-06 | d8 | blood | PB | 2 | 4845 | 4391 | 6590 | 5772 | 11244.5 | 2279 |
| 397-06 | d15 | LN | NS | 1 | 254 | 225 | 6975 | 6905 | 3362 | 1346 |
| 397-06 | d15 | LN | NS | 2 | 240 | 222 | 6086 | 6036 | 3286.5 | 1326.5 |
| 397-06 | d57 | LN | NS | 1 | 118 | 107 | 3408 | 3234 | 3156 | 1291 |
| 397-06 | d57 | LN | NS | 2 | 98 | 85 | 2438 | 2312 | 3253.5 | 1309.5 |
| 397-06 | d121 | LN | NS | 1 | 234 | 218 | 3678 | 3088 | 3378.5 | 1394 |
| 397-06 | d121 | LN | NS | 2 | 270 | 251 | 3379 | 2886 | 3306.5 | 1376 |
| 397-06 | d180 | blood | IgDlo | 1 | 8581 | 7559 | 8634 | 8567 | 4956 | 1665 |
| 397-06 | d180 | blood | IgDlo | 2 | 7185 | 6339 | 7388 | 7329 | 4853 | 1626 |
| 397-06 | d181 | LN | NS | 1 | 349 | 313 | 7271 | 6899 | 3616 | 1421 |
| 397-06 | d181 | LN | NS | 2 | 448 | 414 | 8281 | 7822 | 3724 | 1430 |

Processing of BCR and 5' gene expression data from scRNA-seq of WU397 participants 397-05
and 397-06. LN, lymph node; NS, no sorting.

**Supplementary Table 2.**

| Participant | Time Point | Tissue | Sorting | Replicate | BCR |  | 5' gene expression |  |  |  |
| --- | --- | --- | --- | --- | --- | --- | --- | --- | --- | --- |
|  |  |  |  |  | Pre-QC number of cells | Post-QC number of cells | Pre-QC number of cells | Post-QC number of cells | Median number of UMIs per cell | Median number of genes per cell |
| 397-09 | d0 | blood | IgDlo | 1 | 10857 | 7697 | 10514 | 10429 | 4429 | 1542 |
| 397-09 | d0 | blood | IgDlo | 2 | 13440 | 9494 | 13753 | 13702 | 4209 | 1507 |
| 397-09 | d0 | LN | NS | 1 | 707 | 633 | 4575 | 3858 | 4012 | 1605 |
| 397-09 | d0 | LN | NS | 2 | 826 | 755 | 5007 | 4220 | 3829 | 1583 |
| 397-09 | d8 | blood | PB | 1 | 3887 | 3237 | 4584 | 4017 | 8045 | 1904 |
| 397-09 | d8 | blood | PB | 2 | 3970 | 3305 | 4631 | 4154 | 8030 | 1902.5 |
| 397-09 | d15 | LN | NS | 1 | 601 | 527 | 2176 | 2070 | 3393 | 1328 |
| 397-09 | d15 | LN | NS | 2 | 616 | 555 | 2217 | 2138 | 3260 | 1296 |
| 397-09 | d57 | LN | NS | 1 | 1067 | 953 | 6250 | 6181 | 3746 | 1442 |
| 397-09 | d57 | LN | NS | 2 | 1017 | 910 | 6688 | 6616 | 3741.5 | 1455 |
| 397-09 | d121 | blood | IgDlo | 1 | 7293 | 5456 | 7212 | 7083 | 3901 | 1400 |
| 397-09 | d121 | blood | IgDlo | 2 | 7900 | 6011 | 7624 | 7494 | 3849 | 1406 |
| 397-09 | d121 | LN | NS | 1 | 1243 | 1141 | 4802 | 4643 | 4016 | 1594 |
| 397-09 | d121 | LN | NS | 2 | 1398 | 1286 | 5424 | 5251 | 3655 | 1516 |
| 397-17 | d0 | blood | IgDlo | 1 | 4320 | 3884 | 4456 | 4216 | 4137.5 | 1485 |
| 397-17 | d0 | blood | IgDlo | 2 | 4030 | 3490 | 4252 | 4044 | 4164 | 1472 |
| 397-17 | d0 | LN | NS | 1 | 1799 | 1683 | 5707 | 5642 | 3561 | 1455 |
| 397-17 | d0 | LN | NS | 2 | 1959 | 1823 | 6095 | 6028 | 3542.5 | 1442 |
| 397-17 | d8 | blood | PB | 1 | 3305 | 3060 | 6676 | 3460 | 15321 | 2474 |
| 397-17 | d8 | blood | PB | 2 | 4561 | 3447 | 6157 | 3576 | 11379.5 | 2180.5 |
| 397-17 | d15 | LN | NS | 1 | 2458 | 2274 | 14245 | 13624 | 3330 | 1371.5 |
| 397-17 | d15 | LN | NS | 2 | 2849 | 2600 | 15818 | 14963 | 3283 | 1364 |
| 397-17 | d57 | LN | NS | 1 | 3062 | 2847 | 13994 | 10534 | 3710 | 1489 |
| 397-17 | d57 | LN | NS | 2 | 3534 | 3277 | 14734 | 10845 | 3713 | 1474 |
| 397-17 | d121 | blood | IgDlo | 1 | 3381 | 2677 | 6488 | 4546 | 5141.5 | 1646 |
| 397-17 | d121 | blood | IgDlo | 2 | 5658 | 4010 | 8870 | 6265 | 4747 | 1531 |
| 397-17 | d121 | LN | NS | 1 | 1925 | 1739 | 9508 | 8736 | 3255.5 | 1304 |
| 397-17 | d121 | LN | NS | 2 | 1917 | 1686 | 9765 | 8911 | 3257 | 1316 |
| 397-17 | d181 | LN | NS | 1 | 2061 | 1960 | 10632 | 10529 | 3509 | 1438 |
| 397-17 | d181 | LN | NS | 2 | 2032 | 1933 | 10799 | 10701 | 3509 | 1472 |

Processing of BCR and 5' gene expression data from scRNA-seq of WU397 participants 397-09 and 397-17. LN, lymph node; NS, no sorting.

**Supplementary Table 3.**

| Participant | Time Point | Tissue | Sorting | Replicate | BCR |  | 5' gene expression |  |  |  |
| --- | --- | --- | --- | --- | --- | --- | --- | --- | --- | --- |
|  |  |  |  |  | Pre-QC number of cells | Post-QC number of cells | Pre-QC number of cells | Post-QC number of cells | Median number of UMIs per cell | Median number of genes per cell |
| 397-22 | d0 | blood | IgDlo | 1 | 6467 | 4516 | 5535 | 5356 | 4767.5 | 1573 |
| 397-22 | d0 | blood | IgDlo | 2 | 7884 | 5152 | 6670 | 6508 | 4556.5 | 1505 |
| 397-22 | d0 | LN | NS | 1 | 697 | 618 | 4624 | 4335 | 4304 | 1578 |
| 397-22 | d0 | LN | NS | 2 | 934 | 797 | 4790 | 4526 | 3916.5 | 1443.5 |
| 397-22 | d8 | blood | PB | 1 | 3647 | 3047 | 3854 | 3528 | 10356.5 | 2095.5 |
| 397-22 | d8 | blood | PB | 2 | 2870 | 2440 | 3284 | 3012 | 9889 | 2027 |
| 397-22 | d15 | LN | NS | 1 | 890 | 808 | 5045 | 4965 | 4374 | 1613 |
| 397-22 | d15 | LN | NS | 2 | 1369 | 1248 | 6101 | 6015 | 4320 | 1574 |
| 397-22 | d57 | LN | NS | 1 | 103 | 96 | 5447 | 2947 | 4547 | 1684 |
| 397-22 | d57 | LN | NS | 2 | 92 | 80 | 6373 | 3481 | 4530 | 1645 |
| 397-22 | d121 | LN | NS | 1 | 727 | 682 | 4897 | 4619 | 4453 | 1680 |
| 397-22 | d121 | LN | NS | 2 | 922 | 858 | 6007 | 5720 | 4292.5 | 1590 |
| 397-22 | d180 | blood | IgDlo | 2 | 5589 | 3896 | 6311 | 6061 | 3292 | 1136 |
| 397-22 | d181 | LN | NS | 1 | 997 | 927 | 3908 | 3447 | 4016 | 1430 |
| 397-22 | d181 | LN | NS | 2 | 1269 | 1177 | 4824 | 4315 | 3959 | 1400 |
| 397-28 | d0 | blood | IgDlo | 1 | 4785 | 4343 | 5362 | 4856 | 4687.5 | 1569 |
| 397-28 | d0 | blood | IgDlo | 2 | 5990 | 5367 | 6760 | 6174 | 4952.5 | 1584.5 |
| 397-28 | d0 | LN | NS | 1 | 1687 | 1605 | 6908 | 6884 | 2645 | 1124 |
| 397-28 | d0 | LN | NS | 2 | 1058 | 1015 | 4342 | 4325 | 2598 | 1075 |
| 397-28 | d8 | blood | PB | 1 | 1629 | 1328 | 2138 | 1178 | 14806 | 2599 |
| 397-28 | d8 | blood | PB | 2 | 1978 | 1615 | 2869 | 1636 | 13439 | 2407 |
| 397-28 | d15 | LN | NS | 1 | 2704 | 2579 | 6788 | 6620 | 3688.5 | 1401 |
| 397-28 | d15 | LN | NS | 2 | 2785 | 2653 | 6568 | 6421 | 3735 | 1367 |
| 397-28 | d57 | LN | NS | 1 | 2665 | 2399 | 8524 | 7426 | 3503 | 1316 |
| 397-28 | d57 | LN | NS | 2 | 2454 | 1846 | 9418 | 8260 | 3300 | 1281 |
| 397-28 | d121 | LN | NS | 1 | 306 | 285 | 2371 | 2065 | 3112 | 1253 |
| 397-28 | d121 | LN | NS | 2 | 425 | 410 | 3284 | 2924 | 3062 | 1206 |
| 397-28 | d180 | blood | IgDlo | 1 | 5668 | 4946 | 6199 | 5872 | 4320 | 1358.5 |
| 397-28 | d180 | blood | IgDlo | 2 | 6389 | 5285 | 7233 | 6774 | 4712 | 1433 |
| 397-28 | d181 | LN | NS | 1 | 2509 | 2397 | 13723 | 13699 | 3561 | 1340 |
| 397-28 | d181 | LN | NS | 2 | 2240 | 2133 | 13580 | 13546 | 3726.5 | 1342 |

Processing of BCR and 5' gene expression data from scRNA-seq of WU397 participants 397-22
and 397-28. LN, lymph node; NS, no sorting.

**Supplementary Table 4.**

| Participant | Time Point | Tissue | Sorting | Replicate | BCR |  | 5' gene expression |  |  |  |
| --- | --- | --- | --- | --- | --- | --- | --- | --- | --- | --- |
|  |  |  |  |  | Pre-QC number of cells | Post-QC number of cells | Pre-QC number of cells | Post-QC number of cells | Median number of UMIs per cell | Median number of genes per cell |
| 397-29 | d0 | blood | IgDlo | 1 | 961 | 862 | 929 | 842 | 4791 | 1481.5 |
| 397-29 | d0 | blood | IgDlo | 2 | 871 | 786 | 762 | 729 | 4289 | 1374 |
| 397-29 | d0 | LN | NS | 1 | 1776 | 1660 | 7865 | 7772 | 3448 | 1297 |
| 397-29 | d0 | LN | NS | 2 | 1762 | 1625 | 7593 | 7495 | 3201 | 1235 |
| 397-29 | d8 | blood | PB | 1 | 3499 | 3156 | 6950 | 4072 | 14935 | 2309 |
| 397-29 | d8 | blood | PB | 2 | 3472 | 2950 | 6983 | 3645 | 14930 | 2322 |
| 397-29 | d15 | LN | NS | 1 | 973 | 907 | 6559 | 6508 | 3179 | 1248.5 |
| 397-29 | d15 | LN | NS | 2 | 1101 | 1043 | 8649 | 8593 | 3501 | 1330 |
| 397-29 | d57 | LN | NS | 1 | 2340 | 2126 | 8788 | 8496 | 3866 | 1385 |
| 397-29 | d57 | LN | NS | 2 | 2515 | 2251 | 9449 | 9127 | 3894 | 1383 |
| 397-29 | d121 | LN | NS | 1 | 1879 | 1751 | 7401 | 7276 | 3466.5 | 1269 |
| 397-29 | d121 | LN | NS | 2 | 1951 | 1809 | 7959 | 7802 | 4042.5 | 1423 |
| 397-29 | d180 | blood | IgDlo | 1 | 761 | 667 | 724 | 667 | 4136 | 1315 |
| 397-29 | d180 | blood | IgDlo | 2 | 735 | 654 | 677 | 624 | 4264 | 1378 |
| 397-29 | d181 | LN | NS | 1 | 5931 | 4849 | 9006 | 8925 | 3602 | 1420 |
| 397-29 | d181 | LN | NS | 2 | 6181 | 5456 | 11286 | 11194 | 3618 | 1397 |

Processing of BCR and 5' gene expression data from scRNA-seq of WU397 participant 397-29.
LN, lymph node; NS, no sorting.

**Supplementary Table 5.**
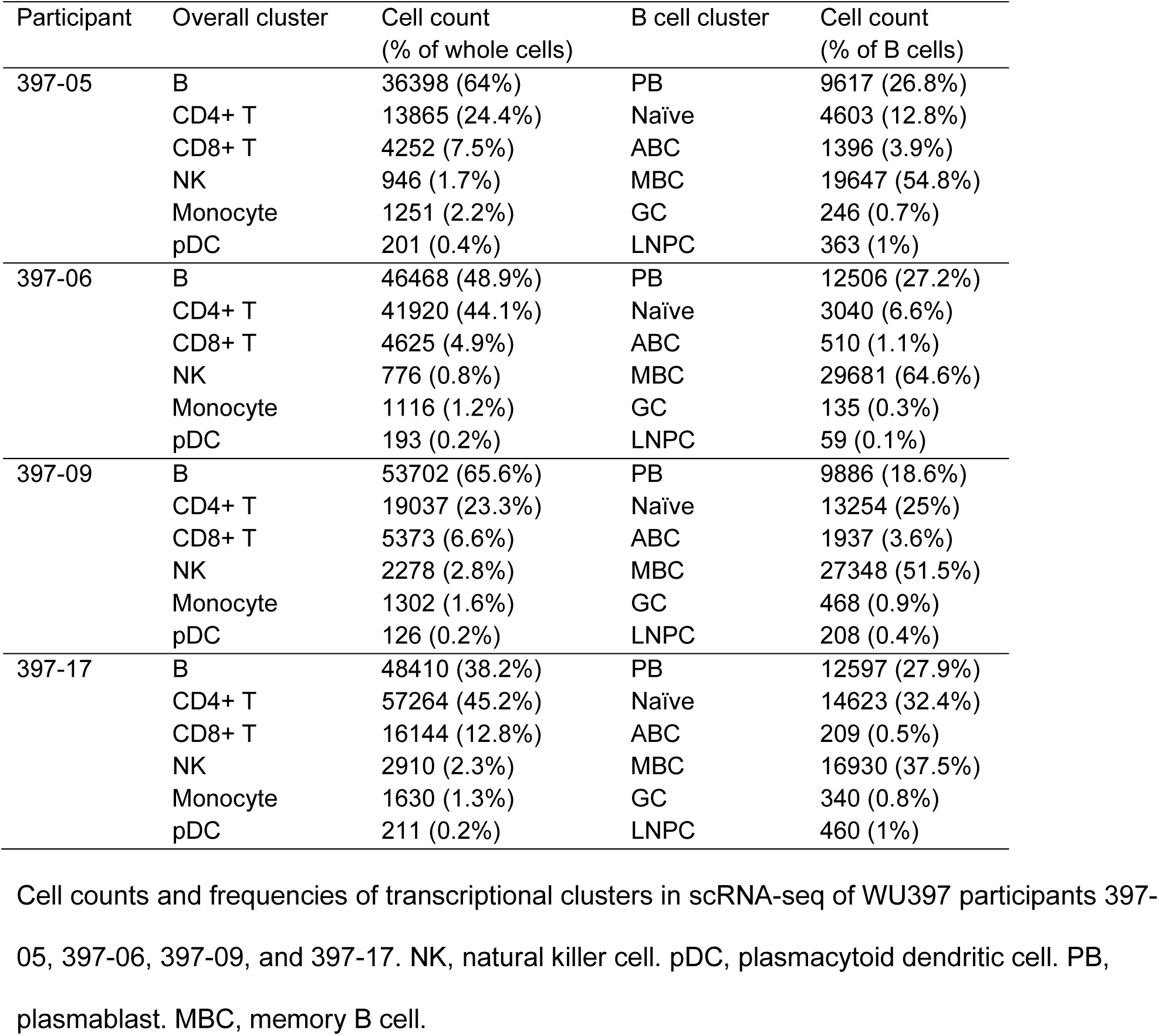

**Supplementary Table 6.**

| Participant | Overall cluster | Cell count<br>(% of whole cells) | B cell cluster | Cell count<br>(% of B cells) |
| --- | --- | --- | --- | --- |
| 397-22 | B | 30742 (44.7%) | PB | 7851 (26.2%) |
|  | CD4+ T | 25357 (36.9%) | Naïve | 2228 (7.4%) |
|  | CD8+ T | 9612 (14%) | ABC | 902 (3%) |
|  | NK | 1526 (2.2%) | MBC | 17484 (58.5%) |
|  | Monocyte | 1275 (1.9%) | GC | 1169 (3.9%) |
|  | pDC | 290 (0.4%) | LNPC | 278 (0.9%) |
| 397-28 | B | 45430 (46.1%) | PB | 3097 (7.3%) |
|  | CD4+ T | 42651 (43.3%) | Naïve | 17462 (41.1%) |
|  | CD8+ T | 7516 (7.6%) | ABC | 338 (0.8%) |
|  | NK | 1239 (1.3%) | MBC | 19962 (47%) |
|  | Monocyte | 1425 (1.4%) | GC | 1148 (2.7%) |
|  | pDC | 344 (0.3%) | LNPC | 487 (1.1%) |
| 397-29 | B | 32166 (34.3%) | PB | 6039 (21.5%) |
|  | CD4+ T | 50195 (53.6%) | Naïve | 4412 (15.7%) |
|  | CD8+ T | 8102 (8.6%) | ABC | 222 (0.8%) |
|  | NK | 1195 (1.3%) | MBC | 13509 (48.2%) |
|  | Monocyte | 1511 (1.6%) | GC | 3302 (11.8%) |
|  | pDC | 520 (0.6%) | LNPC | 540 (1.9%) |

Cell counts and frequencies of transcriptional clusters in scRNA-seq of WU397 participants 397- 22, 397-28, and 397-29. NK, natural killer cell. pDC, plasmacytoid dendritic cell. PB, plasmablast. MBC, memory B cell.

## Methods

### Human subjects and study design

Study WU397 was approved by the Institutional Review Board of Washington University in St Louis (IRB 2208058) and written consent was obtained from all participants. Twenty-nine participants were enrolled. Participants were aged 23-51 years old. Participants reported no adverse effects. No statistical methods were used to predetermine sample size. Investigators were not blinded to experiments and outcome assessment. Blood samples were obtained by standard phlebotomy. PBMCs were isolated using Vacutainer CPT tubes (BD); the remaining red blood cells were lysed with ammonium chloride lysis buffer (Lonza). Cells were immediately used or cryopreserved in 10% dimethylsulfoxide (DMSO) in fetal bovine serum (FBS). Three attending radiologists with expertise in ultrasound performed the ultrasound guided fine-needle aspiration (FNA) of axillary lymph nodes. Ultrasound was performed of the axilla with a high frequency (11-14 MHz), linear transducer to identify the most accessible lateral axillary node closest to the central axillary vessels. Characteristic features and dimensions of the selected baseline node were documented in annotated images and cine clips, including size, vascularity, distance to central axillary vessels and distance to skin, in order to facilitate identification and FNA of the same lymph node at follow-up. Six passes, each consisting of approximately 60 needle throws, were made using 25-gauge needles, which were flushed with 3 mL of RPMI 1640 supplemented with 10% FBS and 100 U/ml penicillin/streptomycin, followed by three 1 mL rinses. Red blood cells were lysed with ammonium chloride lysis buffer (Lonza). Cells were washed twice with phosphate buffered saline (PBS) supplemented with 2% FBS and 2 mM EDTA and immediately used or cryopreserved in 10% DMSO in FBS.

### Vaccines and antigens

Fluarix quadrivalent influenza vaccine (Northern Hemisphere 2022-2023 season) was purchased from GlaxoSmithKline Biologicals. Seasonal quadrivalent mRNA-1010 vaccine (Northern Hemisphere 2022-2023 season) was provided by Moderna, Inc. For ELISpot, 293F-expressed recombinant HA proteins derived from H1N1 (A/Wisconsin/588/2019), H3N2 (A/Darwin/6/2021), B/Yamagata/16/88-like lineage (B/Phuket/3073/2013), or B/Victoria/2/87-like lineage (B/Austria/1359417/2021) were provided by Moderna, Inc. For ELISA, 293F-expressed recombinant HA proteins were provided by Moderna, Inc. (A/Wisconsin/588/2019, B/Phuket/3073/2013) or purchased from SinoBiological (A/Darwin/6/2021, B/Austria/1359417/2021). For flow cytometry staining, 293F-expressed recombinant HA proteins (A/Wisconsin/588/2019, A/Darwin/6/2021, B/Austria/1359417/2021 HA trimers, B/Phuket/3073/2013 HA1) were purchased from SinoBiological. Recombinant HA was biotinylated using the EZ-Link Micro NHS-PEG4-Biotinylation Kit (Thermo Scientific); excess biotin was removed using 7-kDa Zeba desalting columns (Pierce). For biolayer interferometry (BLI), recombinant 6x His-tagged HA proteins from influenza strains (A/Wisconsin/588/2019, A/Darwin/6/2021, B/Phuket/3073/2013, B/Austria/1359417/2021) expressed in 293F cells were purchased from Immune Technology Corp. For Ig-Seq, recombinant 6x His-tagged HA protein from H3N2 (A/Darwin/6/2021) expressed in 293F cells was purchased from SinoBiological. For multiplex fluorescent bead assay, 293F-expressed recombinant HA proteins from the following influenza strains were purchased from Immune Technology Corp: H3N2 strains, A/Thailand/8/2022, A/Darwin/6/2021, A/Cambodia/e0826360/2020, A/South Australia/34/2019, A/Hong Kong/2671/2019, A/Singapore/INFIMH-16-0019/2016, A/Switzerland/8060/2017, A/Victoria/361/2011, A/Brisbane/10/2007; H1N1 strains, A/Wisconsin/67/2022, A/Sydney/5/2021, A/Wisconsin/588/2019, A/Guangdong-Maonan/SWL1536/2019, A/Michigan/45/2015, A/California/04/2009, A/Puerto_Rico/8/1934. Additionally, 293F-expressed recombinant HA proteins from the following influenza strains were purchased from eEnzyme LLC: H3N2 strains, A/Texas/50/2012, A/Perth/16/2009, A/Hong Kong/1/1968; H1N1 strains, A/Hawaii/70/2019, A/Brisbane/2/2018.

### Viruses

Influenza strains (A/Wisconsin/588/2019, A/Darwin/6/2021, B/Phuket/3073/2013, and B/Austria/1359417/2021) were provided by Dr. Richard Webby (St. Jude Children’s Research Hospital).

### ELISpot

Direct ex-vivo ELISpot was performed to determine the total and vaccine-binding IgG-, IgA-, and IgM-secreting cells in PBMCs. ELISpot plates were coated overnight at 4°C with 3 µg/mL recombinant HA protein and 10 µg/mL anti-human Ig kappa and lambda light chain (Cellular Technology Limited). Secreting cells were detected using three-color FluoroSpot IgA/IgG/IgM ELISpot Kits (Cellular Technology Limited) according to the manufacturer’s instructions. ELISpot plates were analyzed using an ELISpot counter (Cellular Technology Limited).

### ELISA

ELISAs were performed in MaxisSorp 96-well plates (Thermo Fisher Scientific). Wells were coated with 1 µg/mL recombinant HA antigens or bovine serum albumin (BSA) in PBS (100 µL) and incubated at 4°C overnight. Plates were blocked with 0.05% Tween 20 and 10% FBS in PBS (blocking buffer). Plasma samples were tested at 1:30 starting dilution in blocking buffer, followed by 7 additional threefold serial dilutions. Recombinant mAbs were diluted to 10 or 2 μg/ml in blocking buffer and added to the plates. Plates were incubated for 90 min at room temperature followed by three washes with 0.05% Tween 20 in PBS. Goat anti-human IgG-HRP (Jackson ImmunoResearch) was diluted 1:2500 in blocking buffer and added to plates. Plates were incubated for 90 min at room temperature followed by three washes with 0.05% Tween 20 in PBS and three washes in PBS. Peroxidase substrate (SigmaFAST o-Phenylenediamine dihydrochloride, Sigma-Aldrich) was used to develop plates. Reactions were stopped by the addition of 1 M HCl. Optical density measurements were taken at 490 nm. The half-maximal binding dilution for plasma was calculated using nonlinear regression (GraphPad Prism v10). The threshold of positivity for recombinant mAbs was set as three times the optical density of background binding to BSA. Recombinant mAbs that demonstrated low cross reactivity at 10 μg/ml were further tested at 2 μg/ml. For testing the avidity of mAbs, ELISAs were performed as above, but wells were coated with 0.1 µg/mL recombinant HA antigens (100 µL). Recombinant mAbs were diluted to 30 µg/mL in blocking buffer, followed by 7 additional threefold serial dilutions. Area under the curve was calculated using GraphPad Prism v10.

### Flow cytometry and cell sorting

For MBC analysis from WU397 participants, cryo-preserved PBMCs were thawed and incubated for 30 min on ice with biotinylated recombinant HA proteins, purified CD16 (3G8, BioLegend, 1:100), CD32 (FUN-2, BioLegend, 1:100), CD64 (10.1, BioLegend, 1:100) in 2% FBS and 2 mM EDTA in PBS (P2). Cells were washed twice, then stained for 30 min on ice with CD38–BB700 (HIT2, BD Horizon, 1:500), CD20–Pacific Blue (2H7, 1:400), CD19–BV750 (HIB19, 1:100), IgD–PE (IA6-2, 1:200), streptavidin-BV650 (1:400), CD3–FITC (HIT3a, 1:200), and Zombie NIR (all BioLegend) diluted in Brilliant Staining buffer (BD Horizon). Cells were washed twice with P2. Samples were resuspended in P2 and acquired on an Aurora using SpectroFlo v3.3 (Cytek). Flow cytometry data were analysed using FlowJo v10.1 (Treestar).

For analysis, FNA single cell suspensions were incubated for 30 min on ice with biotinylated recombinant HA proteins, purified CD16 (3G8, BioLegend, 1:100), CD32 (FUN-2, BioLegend, 1:100), CD64 (10.1, BioLegend, 1:100), and PD-1-BB515 (EH12.1, BD Horizon, 1:100) in P2. Cells were washed twice, then stained for 30 min on ice with IgG–BV480 (goat polyclonal, Jackson ImmunoResearch, 1:100), IgA–FITC (M24A, Millipore, 1:500), CD8a–A532 (RPA-T8, Thermo, 1:100), CD38–BB700 (HIT2, BD Horizon, 1:500), CD71-PE-Cy7 (CY1G4, 1:400), CD20–Pacific Blue (2H7, 1:400), CD4–Spark Violet 538 (SK3, 1:400), CD19–BV750 (HIB19, 1:100), IgD–BV785 (IA6-2, 1:200), CXCR5–PE-Dazzle 594 (J252D4, 1:50), CD14–Spark UV 387 (S18004B, 1:100), CD27–PE-Fire810 (O323, 1:200), IgM-BV605 (MHM-88, 1:100), CD3–APC-Fire810 (SK7, 1:50), and Zombie NIR (all BioLegend) diluted in Brilliant Staining buffer (BD Horizon). Cells were washed twice with P2, fixed for 1 h at 25 °C using the True Nuclear fixation kit (BioLegend), washed twice with True Nuclear Permeabilization/Wash buffer, stained with Ki-67-BV711 (Ki-67, BioLegend, 1:200), Blimp1–PE (646702, R&D, 1:100), FOXP3–Spark 685 (206D, BioLegend, 1:200), and Bcl6–R718 (K112-91, BD Horizon, 1:200) for 1 h at 25 °C, and washed twice with True Nuclear Permeabilization/Wash buffer. Samples were resuspended in P2 and acquired on an Aurora using SpectroFlo v3.3 (Cytek). Flow cytometry data were analysed using FlowJo v10.1 (Treestar).

For sorting PBs and MBCs from WU397 participants, cryo-preserved PBMCs collected at baseline (MBCs), 1 week (PBs), and 26 weeks (MBCs) post vaccination were stained for 30 min on ice with CD20–Pacific Blue (2H7, 1:400), IgD–PerCP–Cy5.5 (IA6-2, 1:200), CD19–PE (HIB19, 1:200), CD38–BV605 (HIT2, 1:100), CD3-FITC (HIT3a, 1:200), and Zombie NIR (all BioLegend) diluted in P2. Cells were washed twice, and PBs (live singlet CD3^−^CD19^+^IgD^lo^CD20^lo^CD38^+^) or MBCs (live singlet CD3^−^CD19^+^IgD^lo^) were sorted using a Bigfoot (Invitrogen) into PBS supplemented with 0.05% BSA and immediately processed for scRNA-seq.

### Samples for scRNA-seq

Sorted PBs, sorted MBCs, and lymph node FNA samples were processed using the following 10x Genomics kits: Chromium Next GEM Single Cell 5′ Kit v2 (PN-1000263); Chromium Next GEM Chip K Single Cell Kit (PN-1000286); BCR Amplification Kit (PN-1000253); Dual Index Kit TT Set A (PN-1000215). Chromium Single Cell 5′ Gene Expression Dual Index libraries and Chromium Single Cell V(D)J Dual Index libraries were prepared according to manufacturer’s instructions. Both gene expression and V(D)J libraries were sequenced on a NovaSeq 6000 (Illumina), targeting a median sequencing depth of 50,000 and 5,000 read pairs per cell, respectively.

### Processing of 10x Genomics single-cell BCR reads

Demultiplexed pair-end FASTQ reads were preprocessed using Cell Ranger v.6.0.1 as previously described^7^ (Supplementary Tables 1-4). Initial germline V(D)J gene annotation was performed on the preprocessed BCRs using IgBLAST v.1.18.0^34^ with the deduplicated version of IMGT/V-QUEST reference directory release 202150-3^35^. Isotype annotation was pulled from the ‘c_call’ column in the ‘filtered_contig_annotations.csv’ files outputted by Cell Ranger. Further sequence-level and cell-level quality controls were performed as previously described^7^. Check against potential cross-sample contamination was performed by examining the presence of any pairwise overlap between samples in terms of BCRs with both identical UMIs and identical V(D)J nucleotide sequences. For each group of cells that originated from different samples but whose BCRs had identical UMI and VDJ sequence, there tended to be only one cell that also had corresponding transcriptomic data. As such, for each group, when there was exactly one cell with transcriptomics-based annotation, only that cell was kept; when there was more than one cell, none of the cells was kept. Altogether, 112 cells were removed from the BCR data. Individualized genotypes were inferred based on sequences that passed all quality controls using TIgGER v.1.0.0^36^ and used to finalize V(D)J annotations. Sequences annotated as non-productively rearranged by IgBLAST were removed from further analysis.

### Clonal lineage inference for single-cell BCR data

B cell clonal lineages were inferred on a by-individual basis based on productively rearranged sequences as previously described^7^. Briefly, paired heavy and light chains were first partitioned based on common V and J gene annotations and CDR3 lengths. Within each partition, pairs whose heavy chain CDR3 nucleotide sequences were within 0.15 normalized Hamming distance from each other were clustered as clones. Following clonal inference, full-length clonal consensus germline sequences were reconstructed using Change-O v.1.2.0^37^.

### Single-cell BCR analysis

A B cell clone was considered HA-specific if it contained any sequence corresponding to a recombinant mAb that was synthesized based on the single-cell BCRs and that tested positive for binding via ELISA. Clonal overlap between B cell compartments was visualized using circlize v.0.4.13^38^. SHM frequency was calculated for each heavy chain sequence using SHazaM v.1.1.0^37^ by counting the number of nucleotide mismatches from the germline sequence in the variable segment leading up to the CDR3. Phylogenetic trees for HA-specific B cell clones containing IgA GC B cells were constructed with heavy chains on a by-participant basis using IgPhyML v1.1.3^39^ and the HLP19 model^40^. Trees were visualized using ggtree v3.10.1^41^.

### Processing of 10x Genomics single-cell 5′ gene expression data

Demultiplexed pair-end FASTQ reads were first preprocessed on a by-sample basis and samples involved in a given analysis were subsequently subsampled to the same effective sequencing length and aggregated using Cell Ranger v.6.0.1 as previously described^7^. Quality control was performed on the aggregate gene expression matrix consisting of 681,188 cells and 36,601 features using SCANPY v.1.8.2^42^. Briefly, to remove presumably lysed cells, cells with mitochondrial content greater than 30% of all transcripts were removed. To remove likely doublets, cells with more than 8,000 features or 80,000 total UMIs were removed. To remove cells with no detectable expression of common endogenous genes, cells with no transcript for any of a list of 34 housekeeping genes^7^ were removed. The feature matrix was subset, based on their biotypes, to protein-coding, immunoglobulin, and T cell receptor genes that were expressed in at least 0.05% of the cells in any sample. The resultant feature matrix contained 16,539 genes. Finally, cells with detectable expression of fewer than 200 genes were removed.

The same quality control criteria were applied when samples were analyzed on a by-participant basis for selection of B cells for expression as monoclonal antibodies. All expressed cells were included in the analysis combining all participants, regardless of their quality control metrics in the combined iteration. After quality control, there were a total of 621,877 cells from 109 single-cell samples (Supplementary Tables 5 and 6).

### Single-cell gene expression analysis

Transcriptomic data was analyzed using SCANPY v.1.8.2^42^ as previously described^7^ with minor adjustments suitable for the current datasets. Briefly, overall clusters were first identified using Leiden graph-clustering with resolution 0.12 (Extended Data Figure 3a). UMAPs were faceted by participant and inspected for convergence to assess whether there was a need for integration. Cluster identities were assigned by examining the expression of a set of marker genes^22^ for different cell types (Extended Data Figure 3b). To remove potential contamination by platelets, 383 cells with a log-normalized expression value of >2.5 for *PPBP* were removed. Cells from the overall B cell cluster were further clustered to identify B cell subsets using Leiden graph-clustering resolution 0.95 (Fig 2c and Extended Data Figure 3c, Supplementary Tables 5 and 6). Cluster identities were assigned by examining the expression of a set of marker genes^22^ for different B cell subsets (Extended Data Figure 3d) along with the availability of BCRs. Clusters 6 and 22 were further clustered at resolution 0.10 in order to differentiate naïve, MBC, and PB/LNPC. Cells found in the PB/LNPC clusters that came from blood samples were labelled PB, while those that came from FNA samples were labelled LNPC. Cells found in the GC B cell clusters but which came from blood samples and which had a PB-like expression profile were labelled PB. Although clusters 10 and 17 clustered with B cells during overall clustering, they were labelled “B & T” as their cells tended to have both BCRs and relatively high expression levels of *CD2* and *CD3E*. Cluster 23 showed no marked expression level of any marker gene and was labelled “Unassigned”. The “B & T” and “Unassigned” clusters were subsequently excluded from the final B cell clustering. Heavy chain SHM frequency and isotype usage of the B cell subsets were inspected for consistency with expected values to further confirm their assigned identities.

### Selection of single-cell BCRs for expression

Single-cell gene expression analysis was first performed on a by-participant basis. The number of B cell clones to be expressed was determined by balancing cost and maximizing coverage. For clones found in the week 1 PB compartment but not in the GC B cell or LNPC compartments at any time point, one week 1 PB per clone was selected from every such clone with a clone size of at least 4 cells. For clones found in the GC B cell or LNPC compartments at any time point but not in the week 1 PB compartment, one GC B cell or LNPC per clone was selected from every such clone with a clone size of at least 3 cells from all participants except participant 397-06, and from every such clone from participant 397-06. Additionally, in order for every such clone from participants 397-17 and 397-29 which persisted through week 26 in the GC B cell or LNPC compartments to be expressed, one week 26 GC B cell or LNPC per clone was selected from every such clone from participants 397-17 and 397-29 that had a clone size of 1 or 2 cells and that persisted through week 26 in the GC B cell or LNPC compartments. For clones found in both the week 1 PB compartment and the GC B cell or LNPC compartments at any time point, one week 1 PB or one GC B cell or LNPC per clone was selected from every such clone from all participants except participant 397-05, and from every such clone with a clone size of at least 3 cells from participant 397-05. Lastly, for every HA-specific clone from participants 397-06, 397-17, and 397-29 that was found in both the week 1 PB and the week 26 GC B cell compartments, if not already expressed, one week 1 PB and/or week 26 GC B cell was selected so that every such clone would have one week 1 PB and one week 26 GC B cell expressed.

For selection, where there were multiple choices in terms of compartments and/or time points, a compartment and/or a time point was first randomly selected. Amongst cells with the matching compartment and/or time point, the cell with the highest heavy chain UMI count was then selected, breaking ties based on IGHV SHM frequency. In all selected cells, native pairing was preserved. The selected BCRs were curated as previously described^7^ prior to synthesis.

### Recombinant monoclonal antibodies and fragment antigen binding production

Selected pairs of heavy and light chain sequences were synthesized by GenScript and sequentially cloned into IgG1, Igκ/λ, and fragment antigen binding (Fab) expression vectors. Heavy and light chain plasmids were co-transfected into Expi293F cells (Thermo Fisher Scientific) for recombinant monoclonal antibody production, followed by purification with protein A agarose resin (GoldBio). Expi293F cells were cultured in Expi293 Expression Medium (Gibco) according to the manufacturer’s protocol.

### Hemagglutination inhibition assay

One volume of serum was treated with four volumes of receptor destroying enzyme (RDE) (Seiken, Japan) at 37°C overnight before inactivation at 56 °C for 1 h. In a 96-well U-bottomed plate (Greiner Bio-One, Austria), serial two-fold dilutions of 25 µl RDE treated sera from a 1/2 dilution in phosphate buffered saline (PBS) (resulting in a 1/10 diultion of sera) were incubated with 25 µl (4 hemagglutinating units) of influenza virus in duplicate wells for 1 h. Subsequently, 50 µl of 1.0% (v/v) turkey erythrocytes (Lampire Biological Laboratories, USA) were added to each well. After 30 min incubation, the individual HAI titers were read as the reciprocal of the highest dilution at which 100% hemagglutination was inhibited. The geometric mean HAI titer (GMT) was calculated for each subject and titers < 10 were assigned a value of 5 for calculation purposes. All serum samples were tested for non-specific binding to turkey erythrocytes prior to performing the assay.

### Biolayer interferometry

Kinetic binding studies were performed on an Octet-R8 (Sartorius) instrument. His tags were removed from Fabs using the thrombin cleavage site preceding the tag. Fabs were treated with biotin-tagged thrombin protease (Sigma-Aldrich) for 2 h at room temperature, followed by removal of remaining His-tagged Fabs with HisPur Ni-NTA resin (Thermo Scientific). The thrombin-protease was removed via Streptavidin Sepharose High Performance affinity resin (Cytiva). Anti-Penta-His (HIS1K) sensor tips (Sartorius) were pre-equilibrated in HEPES buffered saline (0.15M sodium chloride, 10 mM HEPES, 3mM EDTA, pH = 7.6) with 0.05% Tween-20 and 1% BSA (kinetic buffer) followed by loading of HA proteins (10 µg/mL) to 0.5 nm. Thrombin-cleaved Fabs diluted in kinetic buffer were loaded for 120 s, then dissociated for 500 s in kinetic buffer. A HIS1K sensor dipping in kinetic buffer was used as reference sensor. Kinetic parameters of reference subtracted kinetic traces were calculated with Octet BLI analysis software v12.1 using a global fit 1:1 binding model.

### H3 binding titer for Ig-seq analysis

EC_50_ values from ELISA were used to determine the H3 A/Darwin/9/2021-specific serum binding titers. First, costar 96-well ELISA plates (Corning, 07-200-721) were coated overnight at 4°C with 4 μg/mL recombinant H3 (SinoBiological, 40859-V08H1) and blocked with a blocking solution containing 1% BSA and 0.05% Tween-20 in PBS (pH 7.4). After blocking, serially diluted serum samples were bound to the plates for 2 hrs at RT, followed by incubation with 1:5000-diluted anti-human IgG Fc secondary antibody-HRP (Invitrogen, 05-4220) for 1 hr. The plate was rinsed with a washing solution containing 0.1% Tween-20 in PBS three times between every step. For detection, 50 μL TMB substrate (Thermo Scientific) was added before quenching with 50 μL 1 M H_2_SO_4_. Absorbance was measured at 450 nm using microplate reader. The EC_50_ values were derived from curve fitting function of GraphPad Prism.

### High-throughput sequencing of V_H_ for Ig-seq analysis

V_H_ amplicon was prepared as previously described^24,43^. Total RNA from PBMCs taken from week 1 post-vaccination sample and reverse transcribed according to the manufacturer’s instructions using SuperScript IV enzyme (Invitrogen) and Oligo(dT) primer (Invitrogen). V_H_ transcripts were amplified using the FastStart High Fidelity PCR System (Roche) with gene-specific primers^44^. V_H_ amplicons were sequenced using the Illumina NextSeq platform. All sequences were annotated and processed using MiXCR 2.1.5^45^.

### Purification of total IgG from serum and subsequent digestion into F(ab′)2

Each plasma sample was first passed through a 3 mL of Protein G agarose (Thermo Fisher, 20397) affinity column in gravity mode. Flow-through was collected and passed through the column three times. The column was washed with 20 mL of PBS prior to elution with 5 mL of 100 mM glycine-HCl, pH 2.7. The eluted solution, containing total IgG, was immediately neutralized with 0.75 mL of 1 M Tris-HCl, pH 8.0. Purified IgG was digested into F(ab’)2 with 25 μg of IdeS per 1 mg of IgG for 5 hrs on a rotator at 37°C and then incubated with Strep-Tactin agarose (IBA-Lifesciences, 2-1206-025) for 1 hr to remove IdeS.

### Antigen-enrichment of F(ab′)2 and mass spectrometry sample preparation

Recombinant H3 (SinoBiological, 40859-V08H1) was immobilized on N-hydroxysuccinimide (NHS)–activated agarose resin (Thermo Fisher, 26197) by overnight rotation at 4°C. The coupled agarose resins were washed with PBS, and unreacted NHS groups were blocked with 1 M Tris-HCl, pH 7.5 for 30 min at RT. The resins were further washed with PBS and packed into a 0.8 mL centrifuge column (Thermo Fisher, 89868). For each sample, F(ab’)2 was incubated with the individual antigen affinity columns for 2 hrs on a rotator at RT. Flow-through was collected, and the column was washed with 5 mL of PBS. H3-specific F(ab’)2 was eluted with 1% (v/v) formic acid in 0.5 mL fractions. Elution fractions were pooled and concentrated under vacuum to a volume of ∼10 μL and neutralized using 2 M NaOH.

The neutralized elution samples and flow-through samples were denatured with 50 μL of 2,2,2-trifluoroethanol (TFE) and 5 μL of 100 mM dithiothreitol (DTT) at 55°C for 1 hr, and then alkylated by incubation with 3 μL of 550 mM iodoacetamide for 30 min at RT in the dark. Alkylation was quenched with 892 μL of 40 mM Tris-HCl, and protein was digested with trypsin (1:30 (w/w) trypsin/protein) for 16 hrs at 37°C. Formic acid was added to 1% (v/v) to quench the digestion, and the sample volume was concentrated to 150 μL under vacuum. Peptides were then purified using C18 spin columns (Thermo Scientific, 89870), washed three times with 0.1% formic acid, and eluted with a 60% acetonitrile and 0.1% formic acid solution. C18 eluate was concentrated under vacuum centrifugation and resuspended in 50 μL in 5% acetonitrile, 0.1% formic acid.

### LC-MS/MS analysis

Samples were analyzed by liquid chromatography-tandem mass spectrometry on an Easy-nLC 1200 (Thermo Fisher Scientific) coupled to an Orbitrap Fusion Tribrid (Thermo Scientific). Peptides were first loaded onto an Acclaim PepMap RSLC NanoTrap column (Dionex; Thermo Scientific) prior to separation on a 75 μm × 15 cm Acclaim PepMap RSLC C18 column (Dionex; Thermo Scientific) using a 1.6%–76% (v/v) acetonitrile gradient over 90 mins at 300 nL/min. Eluting peptides were injected directly into the mass spectrometer using an EASY-Spray source (Thermo Scientific). The instrument was operated in data-dependent mode with parent ion scans (MS1) collected at 120,000 resolution. Monoisotopic precursor selection and charge state screening were enabled. Ions with charge ≥+2 were selected for collision-induced dissociation fragmentation spectrum acquisition (MS2) in the ion trap, with a maximum of 20 MS2 scans per MS1. Dynamic exclusion was active with a 15-s exclusion time for ions selected more than twice in a 30-s window. Each sample was run three times to generate technical replicate datasets.

### MS/MS data analysis

Participant-specific peptide search databases for MS data acquisition were generated using all V_H_ sequences obtained through BCR-seq and scRNA-seq from each participant involved in this study. V_H_ sequences were grouped into clonotypes on the basis of single-linkage hierarchical clustering, with cluster membership requiring ≥90% identity across the CDRH3 amino acid sequence as measured by edit distance as described^25^. These clustered V_H_ sequences were then combined with a database of background proteins, which included a consensus human protein database (Ensembl 73, longest sequence/gene), decoy V_L_ sequences, and a list of common protein contaminants (MaxQuant) to construct the peptide search database.

Peptide spectra were searched against the database using SEQUEST (Proteome Discoverer 2.4; Thermo Scientific). Searches considered fully tryptic peptides only, allowing up to two missed cleavages. A precursor mass tolerance of 5 ppm and fragment mass tolerance of 0.5 Da were used. Modifications of carbamidomethyl cysteine (static), oxidized methionine, and formylated lysine, serine or threonine (dynamic) were selected. High-confidence peptide-spectrum matches (PSMs) were filtered at a false discovery rate of <1% as calculated by Percolator (q-value <0.01, Proteome Discoverer 2.4; Thermo Scientific). Iso/Leu sequence variants were collapsed into single peptide groups. For each scan, PSMs were ranked first by posterior error probability (PEP), then q-value, and finally XCorr. Only unambiguous top-ranked PSMs were kept; scans with multiple top-ranked PSMs (equivalent PEP, q-value, and XCorr) were designated ambiguous identifications and removed. The average mass deviation (AMD) for each peptide was calculated as described^46^. Peptides with AMD >1.7 ppm were removed. Peptide abundance was calculated from the extracted-ion chromatogram (XIC) peak area, as described^25^. For each peptide, a total XIC area was calculated as the sum of all unique peptide XIC areas of associated precursor ions. The average XIC area across replicate injections was calculated for each sample. For each dataset, the eluate and flow-through abundances were compared and peptides with ≥5-fold higher signal in the elution sample were considered to be antigen-specific.

### Clonotype indexing and peptide-to-clonotype mapping

High-confidence peptides identified through MS/MS analysis were mapped to clonotype clusters. Peptides that uniquely mapped to a single clonotype were considered ‘informative’, and clonotypes detected ≥2 PSM were kept for as high-confidence identifications. The abundance of each antibody clonotype was calculated by summing the XIC areas of the informative peptides mapping to ≥4 amino acids of the CDRH3 region. The amount of each clonotype was calculated by multiplying its relative abundance by the serum titer for that sample. Relative amounts were normalized (ranging from 0 to 1) so that the highest amount for each participant is set to 1.

### Diversity index

Representative CDRH3s matched to peptide sequences identified by MS/MS analysis were used to measure the diversity of CDRH3 within the repertoire. To compare the diversity quantitatively, we calculated the effective number of species (^1^*D*) as previously suggested^27,28^: 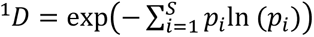, where *p*_i_ is the frequency of the *i* th clone, and S is total number of clones in repertoire.

### B cell lineage analysis for BCR-seq and scRNA-seq

V_H_ sequences obtained through BCR-seq and scRNA-seq were annotated using IgBLAST^34^, and the Immcantation suite pipeline was used to reconstruct the lineage trees^37^. The Immcantation tools and IMGT germline reference were obtained from a pre-packaged Docker container (release version 4.4.0). Briefly, novel V genes were detected with TIgGER^36^, and clonal thresholds were determined using clonal distance. Clonal assignment was performed using hierarchical clustering based on the determined clonal distances. The lineage trees for the representative clonotypes were generated as maximum likelihood lineage tree through IgPhyML with Dowser^40^. Within a lineage tree, each tip is colored based on when its CDRH3 sequence is identified in the proteomics analysis.

### Multiplex fluorescent bead assay

Recombinant HA protein and BSA were incubated for 30 min on ice with different fluorescence intensity peaks of the 7K and 8K Blue Particle Array Kit (Spherotech) at 40 µg/mL, with the exception of recombinant HA A/Perth/16/2009 which was used at 10 µg/mL. Beads were washed twice with 0.05% Tween 20 in PBS, resuspended in plasma samples diluted 1:200 in 0.05% Tween 20 10% FBS in PBS, and incubated for 30 min. Beads were washed twice with 0.05% Tween 20 in PBS, stained with IgG-KIRAVIA Blue 520 (M1310G05, BioLegend, 1:100), incubated for 30 min, washed twice with 0.05% Tween 20 in PBS, and resuspended in 2% FBS and 2 mM EDTA in PBS and acquired on an Aurora using SpectroFlo v3.3 (Cytek). Data were analyzed using FlowJo v10.1 (Treestar). Fold change in median fluorescence intensity was calculated for each week 4 or week 17/26 sample by dividing its median fluorescence intensity by the median fluorescence intensity of the corresponding baseline sample.

### Influenza strains phylogenetic tree

Amino acid sequences for influenza HA proteins were obtained from the GISAID EpiFlu™ Database. Sequences were aligned using Clustal Omega (EMBL-EBI). The resulting sequence alignment was used to generate a phylogenetic tree that was annotated in FigTree v1.4.4 (http://tree.bio.ed.ac.uk/software/figtree/).

## Data Availability

The proteomics data reported in this paper are archived through MassIVE (https://massive.ucsd.edu/ProteoSAFe/static/massive.jsp) under accession code MSV000095155.

## Acknowledgements

The authors thank all study participants for providing samples, members of the Washington University School of Medicine Infectious Disease Clinical Research Unit for WU397 study coordination (study coordinators D. Carani, A. Haile, J. Hajare, R. Thompson, J. Wing; pharmacist M. Royal), the staff of the Center for Clinical Research Imaging at Washington University School of Medicine, P. Rudick, and M. Reiss for assistance with sample collection, and the Genome Technology Access Center in the Department of Genetics at Washington University School of Medicine. The Center is partially supported by NCI Cancer Center Support Grant #P30 CA91842 to the Siteman Cancer Center from the US National Institutes of Health (NIH). The WU397 study was reviewed and approved by the Washington University Institutional Review Board (approval no. 2208058). This work was supported in part with funding from the NIH National Institute of Allergy and Infectious Diseases (NIAID) and Moderna, Inc. The Ellebedy laboratory was supported by NIAID grants U01AI141990 and U01AI144616, and contracts 75N93021C00014 and 75N93019C00051. H.C.M. was supported by NIAID training grant T32AI007172. J.L was supported by NIH grant number P20 GM113132 and P01 AI089618. The content of this manuscript is solely the responsibility of the authors and does not necessarily represent the official view of NIH or NIAID.

## Author contributions

A.H.E. conceived and designed the study. M.K.K. and R.M.P. wrote and maintained the IRB protocol, recruited participants, and coordinated sample collection. H.C.M., F.H., K.D., and H.K.K. processed specimens. B.S.S., M.J.H., and W.D.M. supervised lymph node evaluation prior to FNA and performed FNA. H.C.M. and F.H. performed ELISpot. H.C.M. performed ELISA. A.M. performed HAI assays. H.C.M performed flow cytometry and J.S.T. performed cell sorting. J.Q.Z. analyzed scRNA-seq and BCR repertoire data. A.J.S., S.C.H., F.H., and H.C.M. generated and characterized monoclonal antibodies. H.C.M. performed BLI assays. F.H., J.S.T., and H.C.M. performed multiplex bead assay. T.Y., L.P., and J.L. designed and performed BCR-seq and Ig-seq experiments and analyzed the data. H.C.M., A.H.E., N.H.L., R.N., and R.P. analyzed data. A.H.E. supervised experiments and obtained funding. H.C.M, A.H.E., T.Y., and J.L. composed the manuscript. All authors reviewed and edited the manuscript.

## Competing interests

The Ellebedy laboratory and Infectious Disease Clinical Research Unit received funding under sponsored research agreements from Moderna related to the data presented in the current study. The Ellebedy laboratory received funding from Emergent BioSolutions and AbbVie that are unrelated to the data presented in the current study. A.H.E. has received consulting and speaking fees from InBios International, Fimbrion Therapeutics, RGAX, Mubadala Investment Company, Moderna, Pfizer, GSK, Danaher, Third Rock Ventures, Goldman Sachs and Morgan Stanley and is the founder of ImmuneBio Consulting. A. J. S., J.S.T., and A.H.E. are recipients of a licensing agreement with Abbvie that is unrelated to the data presented in the current study. N.H.L., R.N., and R.P. are employees of and shareholders in Moderna, Inc. The authors declare no other competing interests.

